# A generalization of partial least squares regression and correspondence analysis for categorical and mixed data: An application with the ADNI data

**DOI:** 10.1101/598888

**Authors:** Derek Beaton, ADNI, Gilbert Saporta, Hervé Abdi

**Affiliations:** Rotman Research Institute, Baycrest Health Sciences; ADNI; Conservatoire National des Arts et Metiers; Behavioral and Brain Sciences, The University of Texas at Dallas

**Keywords:** generalized singular value decomposition, latent models, genetics, neuroimaging, heterogeneous data

## Abstract

Current large scale studies of brain and behavior typically involve multiple populations, diverse types of data (e.g., genetics, brain structure, behavior, demographics, or “mutli-omics,” and “deep-phenotyping”) measured on various scales of measurement. To analyze these heterogeneous data sets we need simple but flexible methods able to integrate the inherent properties of these complex data sets. Here we introduce partial least squares-correspondence analysis-regression (PLS-CA-R) a method designed to address these constraints. PLS-CA-R generalizes PLS regression to most data types (e.g., continuous, ordinal, categorical, non-negative values). We also show that PLS-CA-R generalizes many “two-table” multivariate techniques and their respective algorithms, such as various PLS approaches, canonical correlation analysis, and redundancy analysis (a.k.a. reduced rank regression).

## 1 Introduction

Today’s large scale and multi-site studies, such as the UK BioBank (https://www.ukbiobank.ac.uk/) and the Rotterdam study (http://www.erasmus-epidemiology.nl/), collect population level data across numerous types and modalities, (e.g., genetics, neurological, behavioral, clinical, and laboratory measures). Other types of large scale studies—typically those that emphasize diseases and disorders—involve a relatively small of participants but collect a very large number of measures on diverse modalities. Some such studies include the Ontario Neurodegenerative Disease Research Initiative (ONDRI) (Farhan et al. 2016) which includes genetics, multiple types of magnetic resonance brain imaging (Duchesne et al. 2019), a wide array of behavioral, cognitive, clinical, and laboratory batteries, as well as many modalities “between” these types, such as ocular imaging, gait/balance (Montero-Odasso et al. 2017), eye tracking, and neuro-pathology. Though large samples (e.g., UK BioBank) and depth of data (e.g., ONDRI) are necessary to understand typical and disordered samples and populations, few statistical or machine learning approaches exist 1) that can accommodate such large (whether “big” or “wide”), complex, and heterogeneous data sets and 2) that also respect the inherent properties of such data.

In many cases, the analysis of a mixture of data types loses information in part because the data need to be transformed to fit the analytical methods; but this analysis also loses inferential power because the standard assumptions may be inappropriate or incorrect. For example, to analyze categorical and continuous data together, a typical— but inappropriate—strategy is to recode the continuous data into categories such as di-Chotomization, trichotomization, or other (often arbitrary) binning strategies. Furthermore, ordinal and Likert scale data—such as responses on many cognitive, behavioral, clinical, and survey instruments—are often incorrectly treated as metric or continuous values (Bürkner & Vuorre 2019). When it comes to genetic data, such as single nucleotide polymorphims (SNPs), the data are almost always recoded by counting the number of the minor homozygote to give: 0 for the major homozygote (two copies of the most frequent of the two alleles), 1 for the heterozygote, and 2 for the minor homozygote. This {0, 1, 2} recoding of genotypes (1) assumes additive linear effects based on the minor homozygote and (2) is often treated as metric/continuous values (as opposed to categorical or ordinal), even when known effects of risk are neither linear nor additive, such as haplotypic effects (Vormfelde & Brockmöller 2007) nor exclusively based on the minor homozygotes, such as ApoE in Alzheimer’s Disease (Genin et al. 2011). Interestingly other, less restrictive, models (e.g., dominant, recessive, genotypic) perform better (Lettre et al. 2007) than the additive model, but these are rarely used.

Here we introduce partial least squares-correspondence analysis-regression (PLS-CA-R): a latent variable regression modeling approach suited for the analysis of complex data sets. We first show that PLS-CA-R generalizes PLS regression (Wold 1975, Wold et al. 1984, Tenenhaus 1998, Abdi 2010), CA (Greenacre 1984, 2010, Lebart et al. 1984), and PLS-CA (Beaton et al. 2016)—which is the “correlation” companion and basis of PLS-CA-R. We then show that PLS-CA-R is a data-type general PLS regression method based on the generalized singular value decomposition (GSVD) that combines the features of CA (to accommodate multiple data types) with the features of PLS-R (as a regression method generalizing ordinary least squares regression when its assumptions are not met).

We illustrate the multiple uses of PLS-CA-R, with data from the Alzheimer’s Disease Neuroimaging Initiative (ADNI). These data include: diagnosis data (mutually exclusive categories), SNPs (genotypes are categorical), multiple behavioral or clinical instruments (that could be ordinal, categorical, or continuous), and several neuroimaging measures and indices (generally either continuous or non-negative). PLS-CA-R can 1) accommodate different data types in a predictive or fitting framework, 2) regress out (i.e., residualize) effects of mixed and collinear data, and 3) reveal latent variables within a well-established framework.

In addition, we provide a Supplemental section that shows how PLS-CA-R provides the basis for further generalizations. These include different optimization schemes (e.g., covariance as in PLS or correlation as in canonical correlation), transformations for alternate metrics, ridge-like regularization, and three types of PLS algorithms (regression, correlation, and canonical). We also provide an R package and examples at https://github.com/derekbeaton/gpls.

This paper is organized as follows. In Section 2 we introduce PLS-CA-R. Next, in Section 3, we illustrate PLS-CA-R on the TADPOLE challenge (https://tadpole.grand-challenge.org/) and additional genetics data from ADNI across three examples: 1) a simple discriminant example with categorical data, 2) a mixed data example that requires residualization, and 3) a larger example of multiple genetic markers and whole brain tracer uptake (non-negative values). Finally in Section 4 we provide a discussion and conclusions.

## 2 Partial least squares-correspondence analysis-regression

### 2.1 Notation

Bold uppercase letters denote matrices (e.g., **X**), bold lowercase letters denote vectors (e.g., **x)**, and italic lowercase letters denote specific elements (e.g., *x*). Upper case italic letters (e.g., *I*) denote cardinality, size, or length where a lower case italic (e.g., *i*) denotes a specific value of an index. A generic element of matrix **X** would be denoted *x*_*i,j*_. Common letters of varying type faces, for example **X, x**, *x*_*i,j*_ come from the same data structure. A preprocessed or transformed version of a matrix **X** is denoted **Z**_**X**_. The sum of matrix will be denoted with _++_ where, for example, the sum of the matrix **X** is *X*_++_. Vectors are assumed to be column vectors unless otherwise specified. Two matrices side-by-side denotes standard matrix multiplication (e.g., **XY).** The operator ⊙ denotes element-wise (Hadamard) multiplication and ∅ denotes element-wise (Hadamard) division. The matrix **I** denotes the identity matrix, **1** denotes a vector 1’s and **0** is a null matrix (all entries are 0). Superscript ^*T*^ denotes the transpose operation, superscript ^−1^ denotes standard matrix inversion, and superscript ^+^ denotes the Moore-Penrose pseudo-inverse. The diagonal operator, denoted diag{}, transforms a vector into a diagonal matrix, or extracts the diagonal of a matrix to produce a vector.

### 2.2 The Generalized SVD and Correspondence Analysis

Given an *I* × *J* matrix **X**, the singular value decomposition (SVD) decomposes **X** as

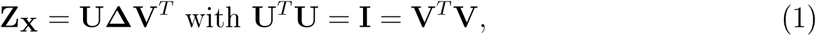

where **X** is of rank *A* (with *A* ≤ min(*I, J*)) and **Δ** is the *A* × *A* diagonal matrix of the singular values, and **Λ** = **Δ**^2^ is the *A* × *A* diagonal matrix of the eigenvalues.

The *generalized* singular value decomposition (GSVD) generalizes the SVD by integrating constraints—also called *metrics* because these matrices define a metric in a generalized Euclidean space—imposed on the rows and columns of the matrix to be decomposed. Specifically, when both an *I* × *I* matrix M and a *J* × *J* matrix **W** are positive definite matrices, the GSVD decomposes **X** as

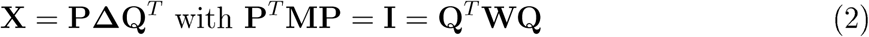

where **X** is of rank *A* and **Δ** is the *A* × *A* diagonal matrix of the singular values, **Λ** = **Δ**^2^ is the *A* × *A* diagonal matrix of the eigenvalues, and **P** and **Q** are the *generalized* singular vectors. We compute singular vectors as 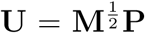 and 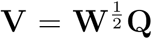 where **U**^*T*^**U** = **I** = **V**^*T*^**V.** From the weights, generalized singular vectors, and singular values we can obtain component (a.k.a. factor) scores as **F**_*I*_ = **MPΔ** and **F**_*J*_ = **WQΔ** for the *I rows* and *J* columns of **X**, respectively. For convenience, the GSVD is presented as a “triplet” with the row metric (weights), data, and column metric (weights) as GSVD(**M, X, W**). We present the GSVD triplet here in slightly different way than it is typically presented (see, e.g., **?**). Our presentation of the GSVD reflects the multiplication steps taken before decomposition (see our Supplmental Material).

Correspondence analysis (CA) is a technique akin to principal components analysis (PCA)—originally designed for the analysis of contingency tables—and analyzes deviations to independence. See Greenacre (1984), Greenacre (2010), and Lebart et al. (1984) for detailed explanations of CA, and then see Escofier-Cordier (1965) and Benzécri (1973) for the origins and early developments of CA. CA analyzes an *I* × *J* matrix **X** whose entries are all non-negative. CA is performed with the GSVD as follows. First the *observed* frequencies matrix is computed as 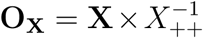. The marginal frequencies from the observed matrix are then computed as

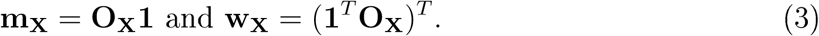

The *expected* frequencies matrix is computed as 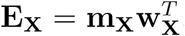. The *deviations* from independence matrix is then

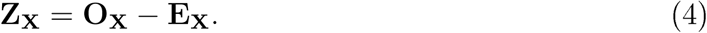

We compute CA from 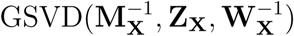 with **M**_**X**_ = diag{**m**_**X**_} and **W**_**X**_ = diag{**w**_**X**_}.

### 2.3 PLS-CA-R

PLS regression (Wold 1975, Wold et al. 1984, 2001) decomposition is an asymmetric and iterative algorithm to find latent variables (a.k.a. components) from a set of predictors from **X** that best explain responses **Y**. PLS-R is asymmetric one data table is privileged (i.e., treated as predictors). PLS-R methods simultaneously decompose **X** and **Y**, generally, through an iterative algorithm. For each iteration, we identify the maximum covariance between **X** mid **Y**, and then deflate each matrix.

Here we present the PLS-CA-R approach which generalizes PLS-R for multiple correspondence analysis (MCA) and correspondence analysis (CA)-like methods that generally apply to categorical (nominal) data. Our formulation of PLS-CA-R provides the basis for generalizations to various data types (e.g., categorical, ordinal, continuous, contingency), as well as different techniques, optimizations, and algorithms (see the Supplemental Material for more details).

For simplicity assume in the following formulation that **X** and **Y** are both complete disjunctive tables as seen in Table 1 (see SEX columns) or Table 2. This formulation also applies generally to non-negative data (see later sections). We define observed matrices for **X** and **Y** as

**Table 1:**
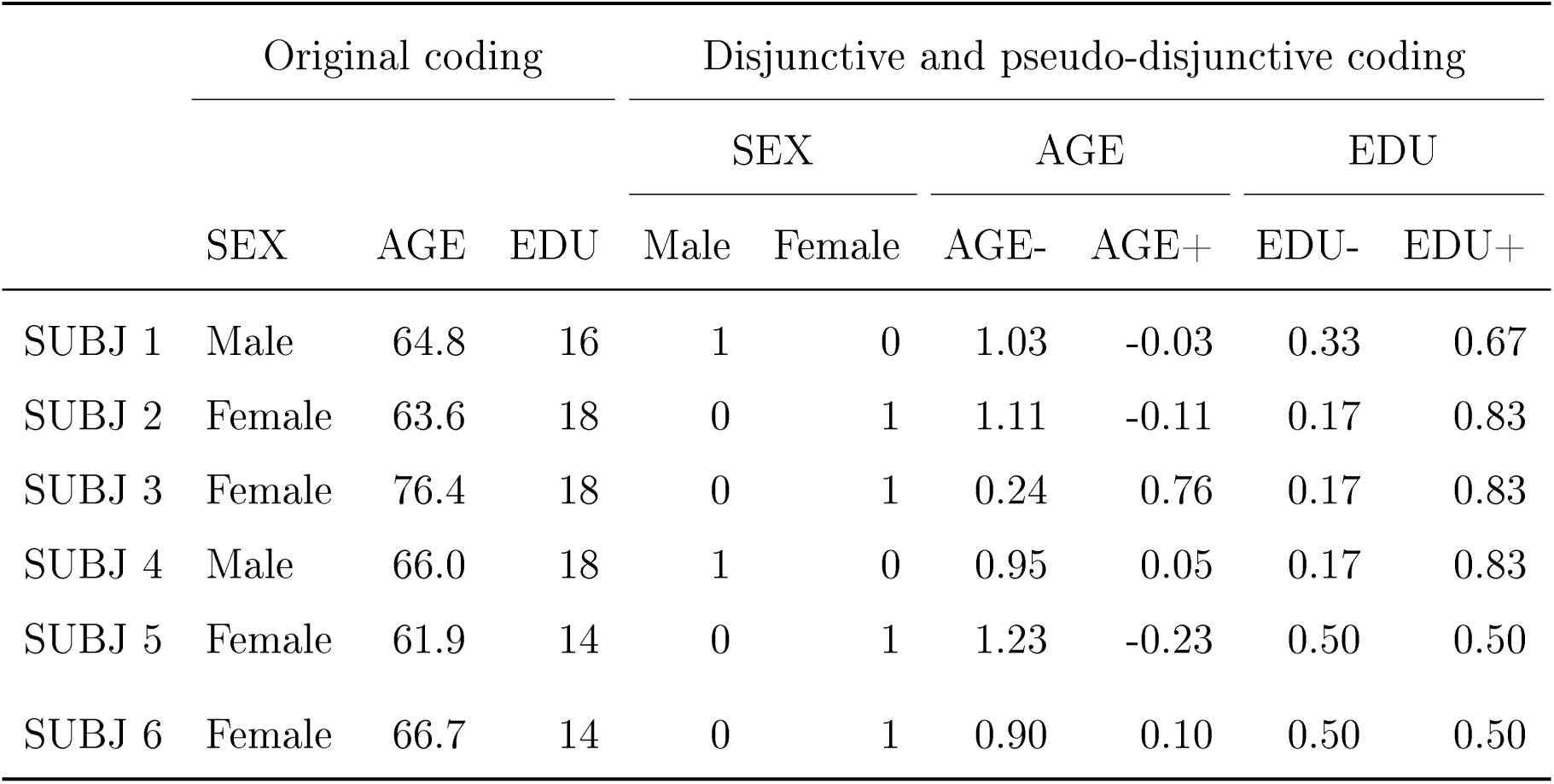
An example of disjunctive (SEX) and pseudo-disjunctive (AGE, EDU) coding through the Escofier or fuzzy transforms. For disjunctive an pseudo-disunctive data, each variable has a row-wise sum of 1 across its respective columns, and thus the row sums across the table are the number of original variables.

**Table 2:**
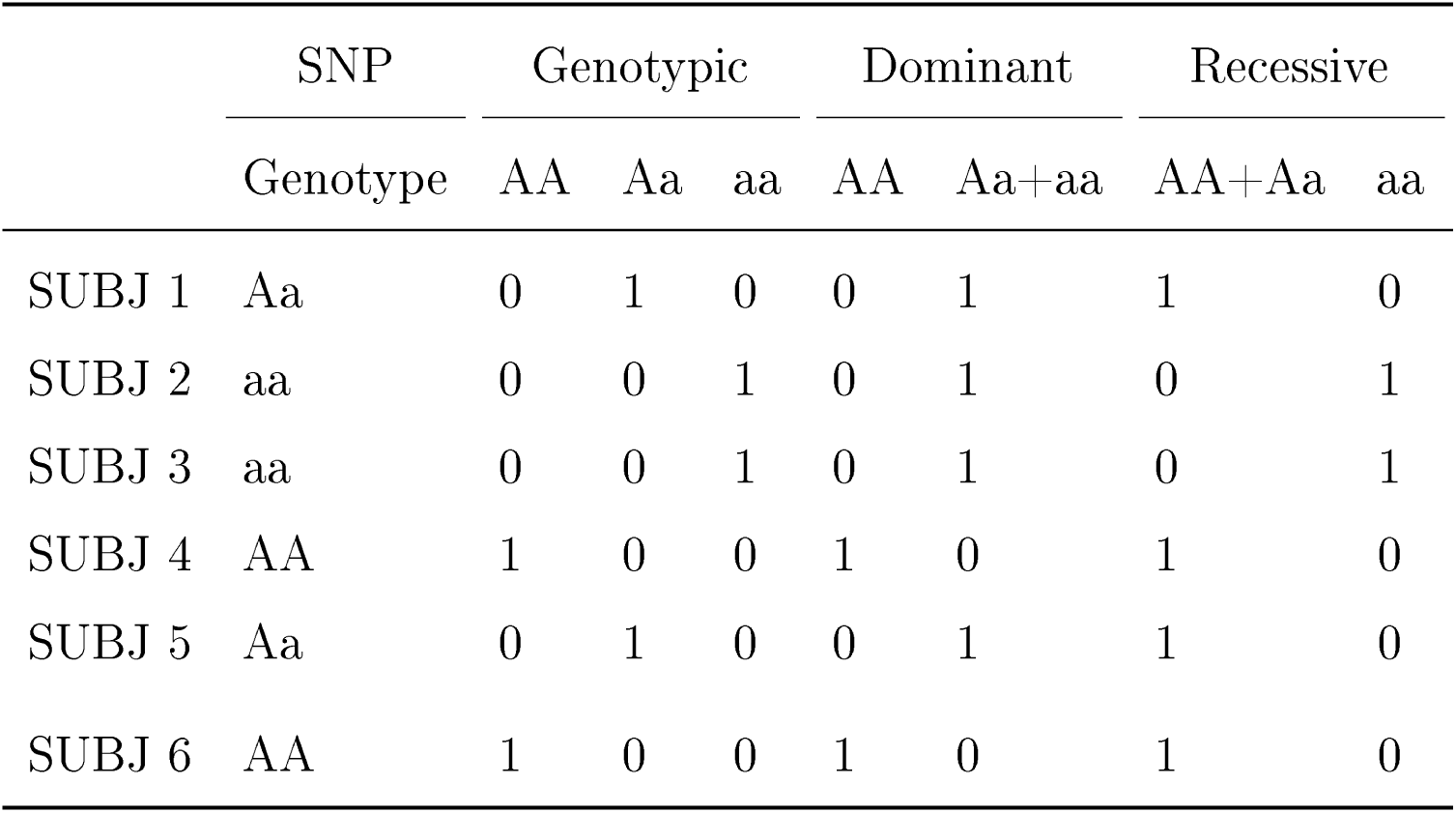
An example of a SNP with its genotypes for respective individuals and disjunctive coding for three types of genetic models: genotypic (three levels), dominant (two levels: major homozygote vs. presence of minor allele), and recessive (two levels: presence of major allele vs. minor homozygote).

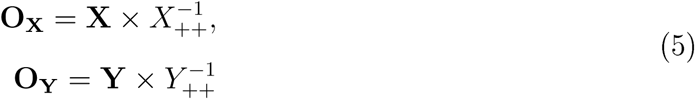

Next we compute marginal frequencies for the rows and columns. We compute row marginal frequencies as

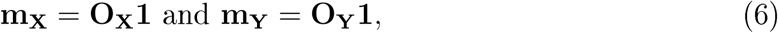

and column frequencies as

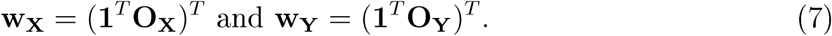

We then define expected matrices as

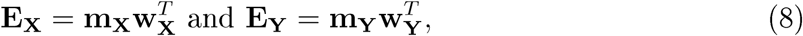

and deviation matrices as

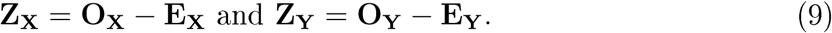

For PLS-CA-R we have row and column weights of **M**_**X**_ = diag{**m**_**X**_}, **M**_**Y**_ = diag{**m**_**Y**_}, **W**_**X**_ = diag{**w**_**X**_}, and **W**_**Y**_ = diag{**w**_**Y**_}. PLS-CA-R makes use of the rank 1 SVD solution iteratively and works as cross-product 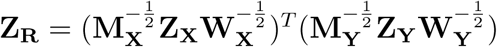, where

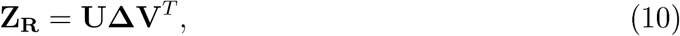

where **U**^*T*^**U** = **I** = **V**^*T*^**V.** We can obtain the *generalized* singular vectors as 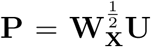 and 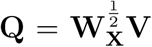, where 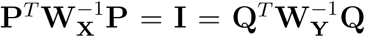. Because we make use of the rank 1 solution iteratively, we only retain the first vectors and values from Eq. 10. We distinguish the retained vectors and values as 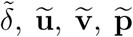, and 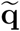. At each iteration, we compute the component scores, for the *J* columns of **Z**_**X**_ and the *K* columns of **Z**_**Y**_, respectively, as 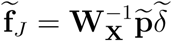 and 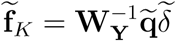. We compute the latent variables as

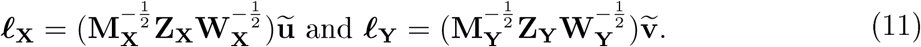

Next we compute 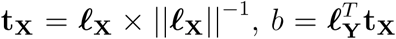, and 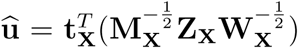. We use **t**_**X**_, b, and **û** to compute rank 1 reconstructed versions of **Z**_**X**_ and **Z**_**Y**_ as

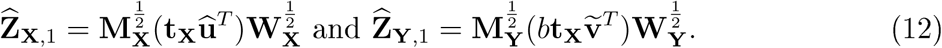

Finally, we deflate **Z**_**X**_ and **Z**_**Y**_ as 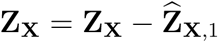 and 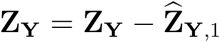. We then repeat the iterative procedure with these deflated **Z**_**X**_ and **Z**_**Y**_. The computations outlined above are performed for *C* iterations where: (1) *C* is some pre-specified number of intended latent variables where *C* ≤ *A* where *A* is the rank of 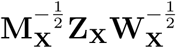 or (2) when **Z**_**X**_ = **0** or **Z**_**Y**_ = **0.** Upon the stopping condition we would have *C* components, and would have collected any vectors into corresponding matrices. Those matrices are:

- two *C* × *C* diagonal mat rices **B** and 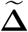 with each *b* and 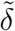 on the diagonal,
- the *I* × *C* matric es **L**_**X**_ **L**_**Y**_, and **T**_**X**_,
- the *J* × *C* matrices 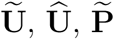, and 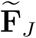, and
- the *K* × *C* matrices 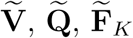.

Algorithm 1 shows the key elements of the PLS-CA-R algorithm, from input to deflation.

### 2.4 Maximization in PLS-CA-R

PLS-CA-R maximizes the common information between **Z**_**X**_ and **Z**_**Y**_ such that

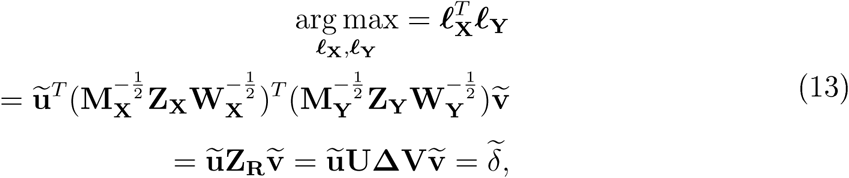

where 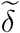 is the first singular value from **Δ** for each *c* step. PLS-CA-R maximization is subject to the orthogonality constraint that 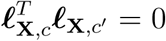 when *c* ≠ *c*′. This orthogonality constraint propagates through to many of the vectors and matrices associated with **Z**_**X**_ where 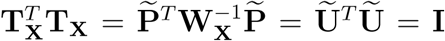 these orthogonality constraints do not apply to the various vectors and matrices associated with **Y**.

#### Algorithm 1: PLS-CA-R algorithm. The results of a rank 1 solution are used to compute the latent variables and values necessary for deflation of Z_X_ and Z>_Y_.

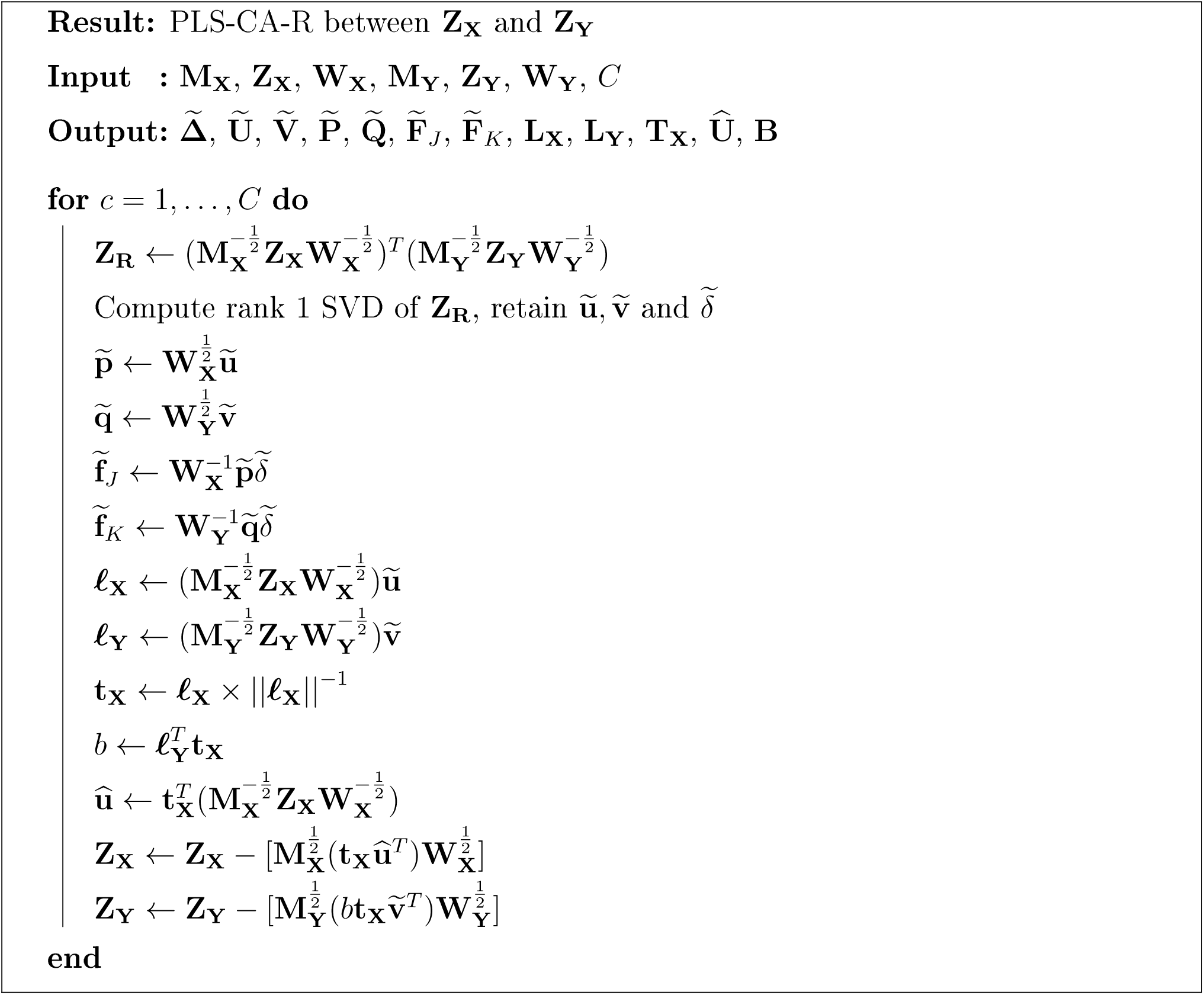

### 2.5 Decomposition and reconstitution

PLS-CA-R is a “double decomposition” where

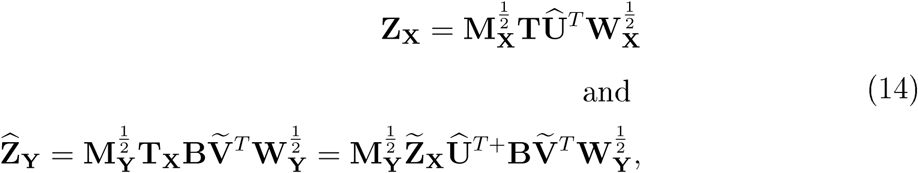

where 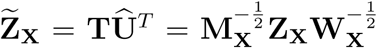. PLS-CA-R, like PLS-R, provides the same values as OLS where

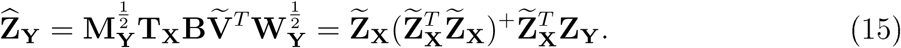

This connection to OLS shows how to residualize (i.e., “regress out” or “correct for”) covariates, akin to how residuals are computed in OLS. We do so with the original **Z**_**Y**_ as 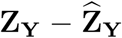. PLS-CA-R produces both a predicted and a residualized version of **Y**. Recall that 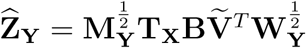. We compute a reconstituted form of **Y** as

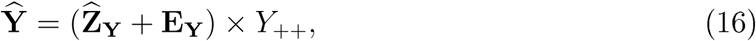

which is the opposite steps of computing the deviations matrix. We add back in the expected values and then scale the data by the total sum of the original matrix. The same can be done for residualized values (i.e., “error”) as

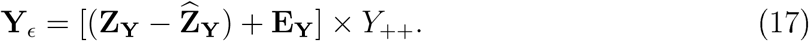

Typically, **E**_**Y**_ is derived the data (as noted in Eq. 8). However, the reconstituted space could come from any model by way of generalized correspondence analysis (GCA) (Escofier 1983, 1984, Grassi & Visentin 1994, Beaton et al. 2018). With GCA we could use any reasonable alternates of **E**_**Y**_ as the model, which could be obtained from, for examples, known priors, a theoretical model, out of sample data, or population estimates. CA can then be applied directly to either **Ŷ** or **Y**_*ϵ*_. The same reconstitution procedures can be applied fitted and residualized versions of **X** as well.

### 2.6 Concluding remarks

Now that we have formalized PLS-CA-R, we want to point out small variations, some caveats, and some additional features. We also briefly discuss here—but show in the Supplemental Material—how PLS-CA-R provides the basis for numerous variations and broader generalizations.

In PLS-CA-R (and PLS-R) each subsequent 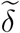 is not guaranteed to be smaller than the previous, with the exception that all 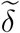 are smaller than the first. This is a by-product of the iterative process and the deflation step. This problem poses two issues: (1) visualization of component scores and (2) explained variance. For visualization of the component scores— which use 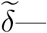there is an alternative computation: 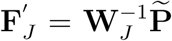 and 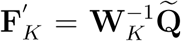. This alternative is referred to as “asymmetric component scores” in the correspondence analysis literature (Abdi & Béra 2018, Greenacre 1993). Additionally, instead of computing the variance per component or latent variable, we can instead compute the amount of variance explained by each component in **X** and **Y**. To do so we require the sum of the eigenvalues of each of the respective matrices per iteration via CA (with the GSVD). Before the first iteration of PLS-CA-R we obtain the full variance (i.e., the sum of the eigenvalues) of each matrix from 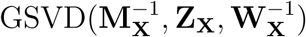 and 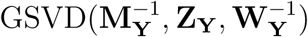, which we respectively refer to as *ϕ*_**X**_ and *ϕ*_**Y**_ We can compute the sum of the eigenvalues for each deflated version of **Z**_**X**_ and **Z**_**Y**_ through the GSVD just as above, referred to as *ϕ*_**X**,*c*_ and *ϕ*_**Y**, *c*_. For each *c* component the proportion of explained variance for each matrix is 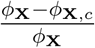 and 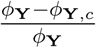.

In our formulation, the weights we use are derived from the χ^2^ assumption of independence. However nearly any choices of weights could be used, so long as the weight matrices are at least positive semi-definite (which requires the use of a generalized inverse). If alternate row weights (i.e., **M**_**X**_ or **M**_**Y**_) were chosen, then the fitted values and residuals are no longer guaranteed to be orthogonal (the same condition is true in weighted OLS).

PLS-CA-R provides an important basis for various extensions. We can use different PLS algorithms in addition to Algorithm 1, for examples, the PLS-SVD and canonical PLS algorithms. Our formulation of PLS-CA-R provides the basis for, and leads to a more generalized approach for other cross-decomposition techniques or optimizations (e.g., canonical correlation, reduced rank regression). This basis also allows for the use of alternate metrics as well as ridge-like regularization. We show how PLS-CA-R leads to these generalizations in the Supplemental Material.

Though we formalized PLS-CA-R as a method for categorical (nominal) data coded in complete disjunctive format (as seen in Table 1—see SEX columns—or Table 2), PLS-CA-R can easily accomodate various data types without loss of information. Specifically, both continuous and ordinal data can be handled with relative ease and in a “pseudo-disjunctive” format, also referred to as “fuzzy coding” where complete disjunctive would be a “crisp coding” (Greenacre 2014). We explain exactly how to handle various data types as Section 3 progresses, which reflects more “real world” problems: complex, mixed data types, and multi-source data.

## 3 Applications & Examples

In this section we provide examples with real data from the Alzheimer’s Disease Neuroimaging Initiative (ADNI). These examples illustrate how to approach mixed data with PLS-CA-R and the multiple uses of PLS-CA-R (e.g., for analyses, as a residualization procedure). We present three sets of analyses. First we introduce of PLS-CA-R through an example where we want to predict genotypes (categorical) from groups (categorical). Next we show how to predict genotypes from a small set Of behavioral and brain variables. This second example illustrates how to: (1) how to recode and analyze mixed data (categorical, ordinal, and continuous), (2) how to use PLS-CA-R, and (3) how to use PLS-CA-R as residualization technique to remove effects from data prior to subsequent analyses. Finally, we present a larger analysis with the goal to predict genotypes from cortical uptake of AV45 (i.e., a radiotracer) PET scan for beta-amyloid (“A*β*”) deposition. This final example also makes use of residualization as illustrated in the second example.

### 3.1 ADNI Data

Data used in the preparation of this article come from the ADNI database (adni.loni.usc.edu). ADNI was launched in 2003 as a public-private funding partnership and includes public funding by the National Institute on Aging, the National Institute of Biomedical Imaging and Bioengineering, and the Food and Drug Administration. The primary goal of ADNI has been to test a wide variety of measures to assess the progression of mild cognitive impairment and early Alzheimer’s disease. The ADNI project is the result of efforts of many coinvestigators from a broad range of academic institutions and private corporations. Michael W. Weiner (VA Medical Center, and University of California-San Francisco) is the ADNI Principal Investigator. Subjects have been recruited from over 50 sites across the United States and Canada (for up-to-date information, see www.adni-info.org).

The data in the following examples come from several modalities from the ADNI-G0/2 cohort. The data come from two sources available from the ADNI download site (http://adni.loni.usc.edu/): genome-wide data and the TADPOLE challenge data (https://tadpole.grand-challenge.org/) which contains a wide variety of data. Because the genetics data are used in every example, we provide all genetics preprocessing details here, and then describe any preprocessing for other data as we discuss specific examples.

For all examples in this paper we use a candidate set of single nucleotide polymorphisms (SNPs) extracted from the genome-wide data. We extracted only SNPs associated with the *MAPT, APP, ApoE*, and *TOMM40* genes because these genes are considered as candidate contributors to various AD pathologies: *MAPT* because it is associated with tau proteins, AD pathology, or cognitive decline (Myers et al. 2005, Trabzuni et al. 2012, Desikan et al. 2015, Cruchaga et al. 2012, Peterson et al. 2014), *APP* because of its association with *β*-amyloid proteins (Cruchaga et al. 2012, Huang et al. 2017, Jonsson et al. 2012), as well as *ApoE* and *TOMM40* because of their strong association with AD pathologies (Linnertz et al. 2014, Roses et al. 2010, Bennet et al. 2010, Huang et al. 2017). SNPs were processed as follows via Purcell et al. (2007) with additional R code as necessary: minor allele frequency (MAF) > 5% and missingness for individuals and genotypes ≤ 10%. Because the SNPs are coded as categorical variables (i.e., for each genotype) we performed an additional level of preprocessing: genotypes > 5% because even with MAF > 5%, it was possible that some genotypes (e.g., the heterozygote or minor homozygote) could still have very few occurrences. Therefore genotypes ≤ 5% were combined with another genotype. In all cases the minor homozygote (^‘^aa’) fell below that threshold and was then combined with its respective heterozygote (^‘^Aa^’^); thus some SNPs were effectively coded as the dominant model (i.e., the major homozygote vs. the presence of a minor allele). See Table for an example of SNP data coding examples. From the ADNI-GO/2 cohort there were 791 available participants with 134 total SNPs across the four candidate genes. These 134 SNPs span 349 columns in disjunctive coding (see Table 2). Other data include diagnosis and demographics, some behavioral and cognitive instruments, and several types of brain-based measures. We discuss these additional data in further detail when we introduce these data.

### 3.2 Diagnosis and genotypes

Our first example asks and answers the question: “which genotypes are associated with which diagnostic category?”. In ADNI, diagnosis at baseline is a categorical variable that denotes which group each participant belongs to (at the first visit): control (CN; *N =* 155), subjective memory complaints (SMC; *N* = 99), early mild cognitive impairment (EMCI; *N* = 277), late mild cognitive impairment (LMCI; *N* = 134), and Alzheimer’s disease (AD; *N* = 126). We present this first example analysis in two ways: akin to a standard regression problem {‘a} la Wold [Wold (1975); Wold et al. (1984); Wold et al. (1987); cf. Eq. 15) and then again in the multivariate perspective of “projection onto latent structures” (Abdi 2010).

For this example we refer to diagnosis groups as the predictors (**X**) and the genotypic data as the responses (**Y**). Both data types are coded in disjunctive format (see Tables 1 and 2). Because there are five columns (groups) in **X**, PLS-CA-R produces only four latent variables (a.k.a. components). Table 4 presents the cumulative explained variance for both **X** and **Y** and shows that groups explain only a small amount of genotypic variance: *R*^2^ = 0.0065.

**Table 3:** Descriptives and demographics for the sample. AD = Alzheimer’s Disease, CN = control, EMCI = early mild cognitive impairment, LMCI = late mild cognitive impairment, SMC = subjective memory complaints.

|  | N | AGE mean (sd) | EDU mean (sd) | Males (Females) |
| --- | --- | --- | --- | --- |
| AD | 126 | 74.53 (8.42) | 15.78 (2.68) | 75 (51) |
| CN | 155 | 74 (6.02) | 16.41 (2.52) | 80 (75) |
| EMCI | 277 | 71.14 (7.39) | 15.93 (2.64) | 156 (121) |
| LMCI | 134 | 72.24 (7.68) | 16.43 (2.57) | 73 (61) |
| SMC | 99 | 72.19 (5.71) | 16.81 (2.51) | 40 (59) |

**Table 4:** The R-squared values over the four latent variables for both groups and genotypes. The full variance of groups is explained over the four latent variables. The groups explained 0.65% of the genotypes.

|  | X (groups) R-squared cumulative | Y (genotypes) R-squared cumulative |
| --- | --- | --- |
| Latent variable 1 | 0.25 | 0.0026 |
| Latent variable 2 | 0.50 | 0.0042 |
| Latent variable 3 | 0.75 | 0.0055 |
| Latent variable 4 | 1.00 | 0.0065 |

**Table 5:** The *a priori* (rows) vs. assigned (columns) accuracies for the discriminant analysis. AD = Alzheimer’s Disease, CN = control, EMCI = early mild cognitive impairment, LMCI = late mild cognitive impairment, SMC = subjective memory complaints.

|  | *Assigned* |  |  |  |  |
| --- | --- | --- | --- | --- | --- |
|  | CN | SMC | EMCI | LMCI | AD |
| CN | 62 | 15 | 54 | 16 | 17 |
| SMC | 20 | 40 | 33 | 18 | 10 |
| EMCI | 34 | 20 | 110 | 23 | 29 |
| LMCI | 14 | 12 | 39 | 44 | 20 |
| AD | 25 | 12 | 41 | 33 | 50 |

In a simple regression-like framework we can compute the variance contributed by genotypes or group (i.e., levels of variables) or variance contributed by entire variables (in this example: SNPs). First we compute the contributions to the variance of the genotypes as the sum of the squared loadings for each item: 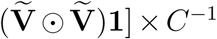, where **1** is a conformable vector of ones. Total contribution takes values between 0 and 1 and describe the proportion of variance for each genotype. Because the contributions are squared loadings, they are additive and so we can compute the contributions for a SNP. A simple criterion to identify genotypes or SNPs that contribute to the model is to identify which genotype or SNP contributes more variance than expected, which is one divided by the total number of original variables (i.e., SNPs). This criterion can be applied on the whole or component-wise. We show the genotypes and SNPs with above expected variance for the whole model (i.e., high contributing variables a regression framework) in Figure 1.

**Figure 1:**
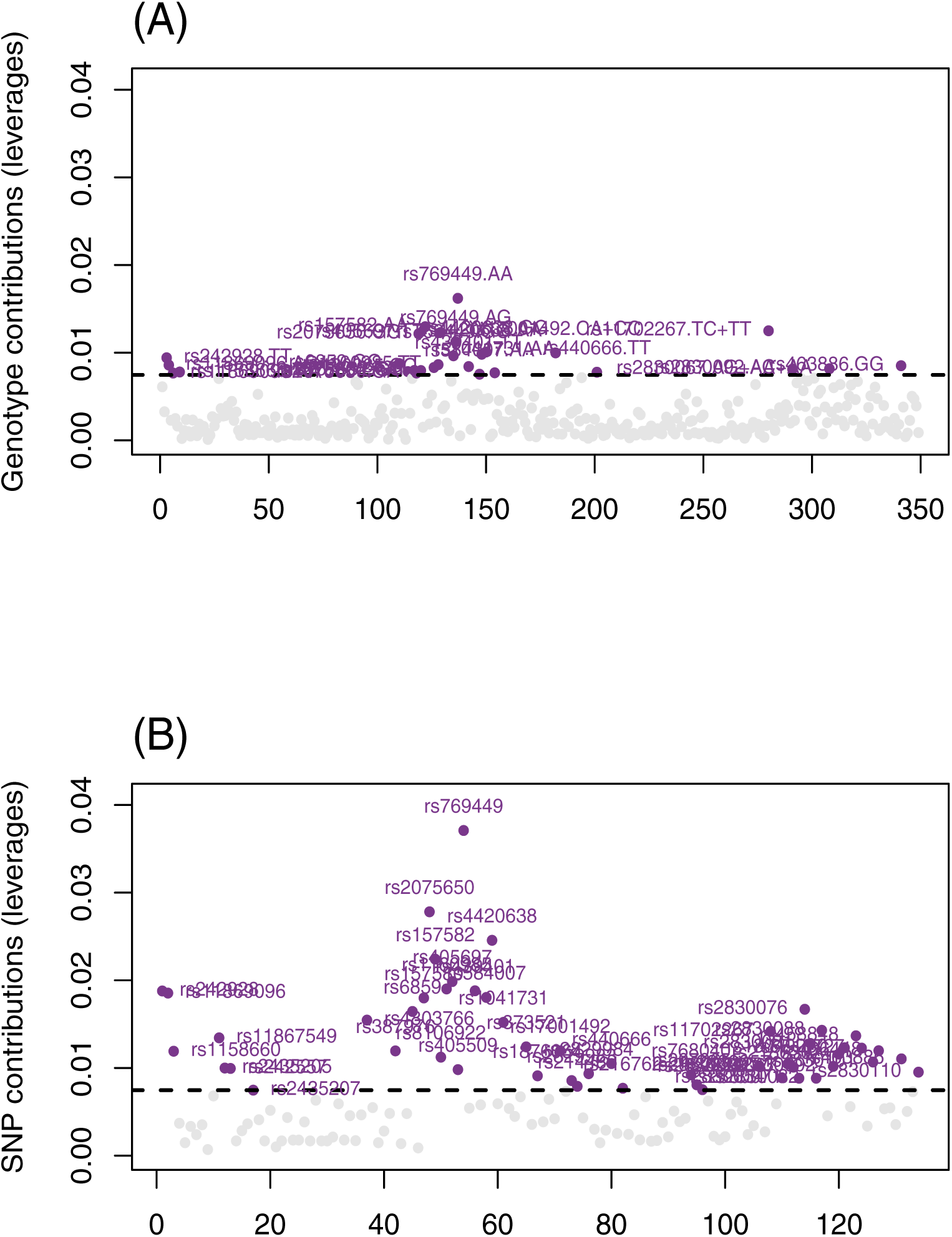
Regression approach to prediction of genotypes from groups. Contributions across all components for genotypes (A; top) and the SNPs (B; bottom) computed as the summation of genotypes within a SNP. The horizontal line shows the expected variance and we only highlight genotypes (A; top) or SNPs (B; bottom) greater than the expected variance. Some of the highest contributing genotypes (e.g., AA and AG genotypes for rs769449) or SNPs (e.g., rs769449 and rs20756560) come from the APOE and TOMM40 genes.

Though PLS-R was initally developed as a regression approach—especially to handle collinear predictors (see explanations in Wold et al. 1984)—it is far more common to use PLS to find latent structures (i.e., components or latent variables) (Abdi 2010). From here on we show the latent variable scores (observations) and component scores (variables) for the first two latent variables/components in Figure 2. The first latent variable (Fig. 2a) shows a gradient from the control (CN) group through to the Alzheimer’s Disease (AD) groups (CN to SMC to EMCI to LMCI to AD). The second latent variable shows a dissociation of the EMCI group from all other groups (Fig. 2b). Figure 2c and d show the component scores for the variables. Genotypes on the left side of first latent variable (horizontal axis in Figs. 2c and d) are more associated with CN and SMC than with the other groups, whereas genotypes on the right side are more associated with AD and LMCI than with the other groups. Through the latent structures approach we can more clearly see the relationships between groups and genotypes. Because we treat the data categorically and code for genotypes, we can identify the specific genotypes that contribute to these effects. For example the ‘AA’ genotype of rs769449 and the ‘GG’ genotype of rs2075650 are more associated with AD and LMCI than with the other groups. In contrast, the ‘TT’ genotype of rs405697 and the ‘TT’ genotype rs439401 are more associated with the CN group than other groups (and thus could suggest potential protective effects).

**Figure 2:**
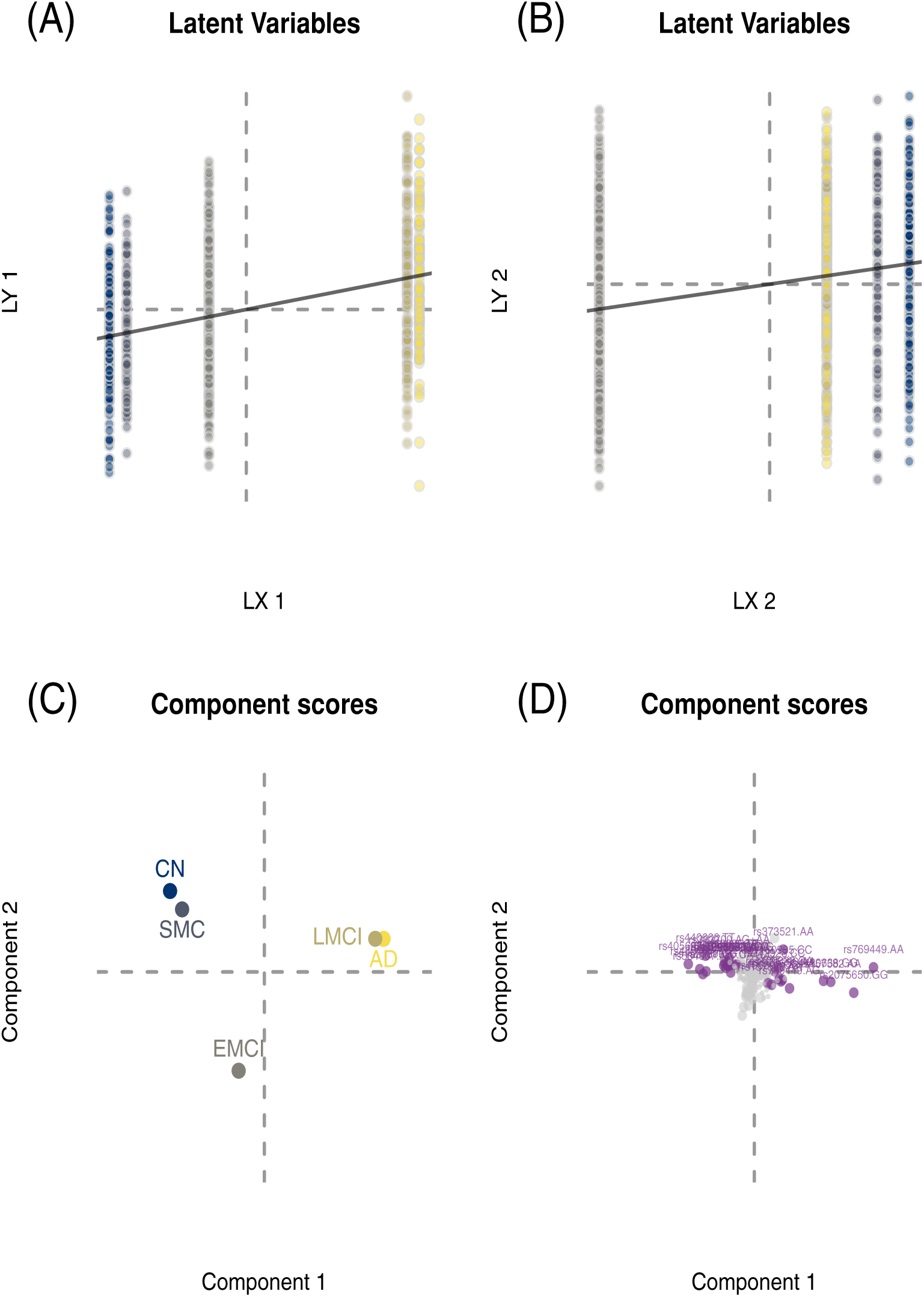
Latent variable projection approach to prediction of genotypes from groups. (A) and (B) show the latent variable scores for latent variables (LVs; components) one and two, respectively; (C) shows the component scores of the groups, and (D) shows the component scores of the genotypes. In (D) we highlight genotypes with above expected contribution to Latent Variable (Component) 1 in purple and make all other genotypes gray.

This group-based analysis is also a discriminant analysis because it maximally separates groups. Thus we can classify observations by assigning them to the closest group. To correctly project observations onto the latent variables we compute **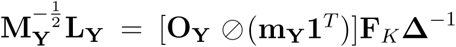** where 1 is a 1 × *K* vector of ones where **O**_**Y**_ ⊘ (**m**_Y_ *1*^*T*^) are “row profiles” of Y (i.e., each element of Y divided by its respective row sum). Observations from **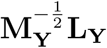**are then assigned to the closest group in **F**_*J*_, either for per component, across a subset of components, or all components. For this example we use the full set (four) of components. The assigned groups can then be compared to the *a priori* groups to compute a classification accuracy. Figure 3 shows the results of the discriminant analysis but only visualized on the first two components. Figures 3a and b respectively show the scores for **F**_*J*_ mid **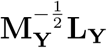** Figure 3c shows the assignment of observations to their closest group. Figure 3d visualizes the accuracy of the assignment, where observations in black are correct assignments (gray are incorrect assignments). The total classification accuracy 38.69% (where chance accuracy was 23.08%). Finally, typical PLS-R discriminant analyses are applied in scenarios where a small set of, or even a single, (typically) categorical responses are predicted from many predictors (Pérez-Enciso & Tenenhaus 2003). However, such an approach appears to be “over optimistic” in its prediction and classification (Rodríguez-Pérez et al. 2018), which is why we present discriminant PLS-CA-R more akin to a typical regression problem (i.e., here a single predictor with multiple responses).

**Figure 3:**
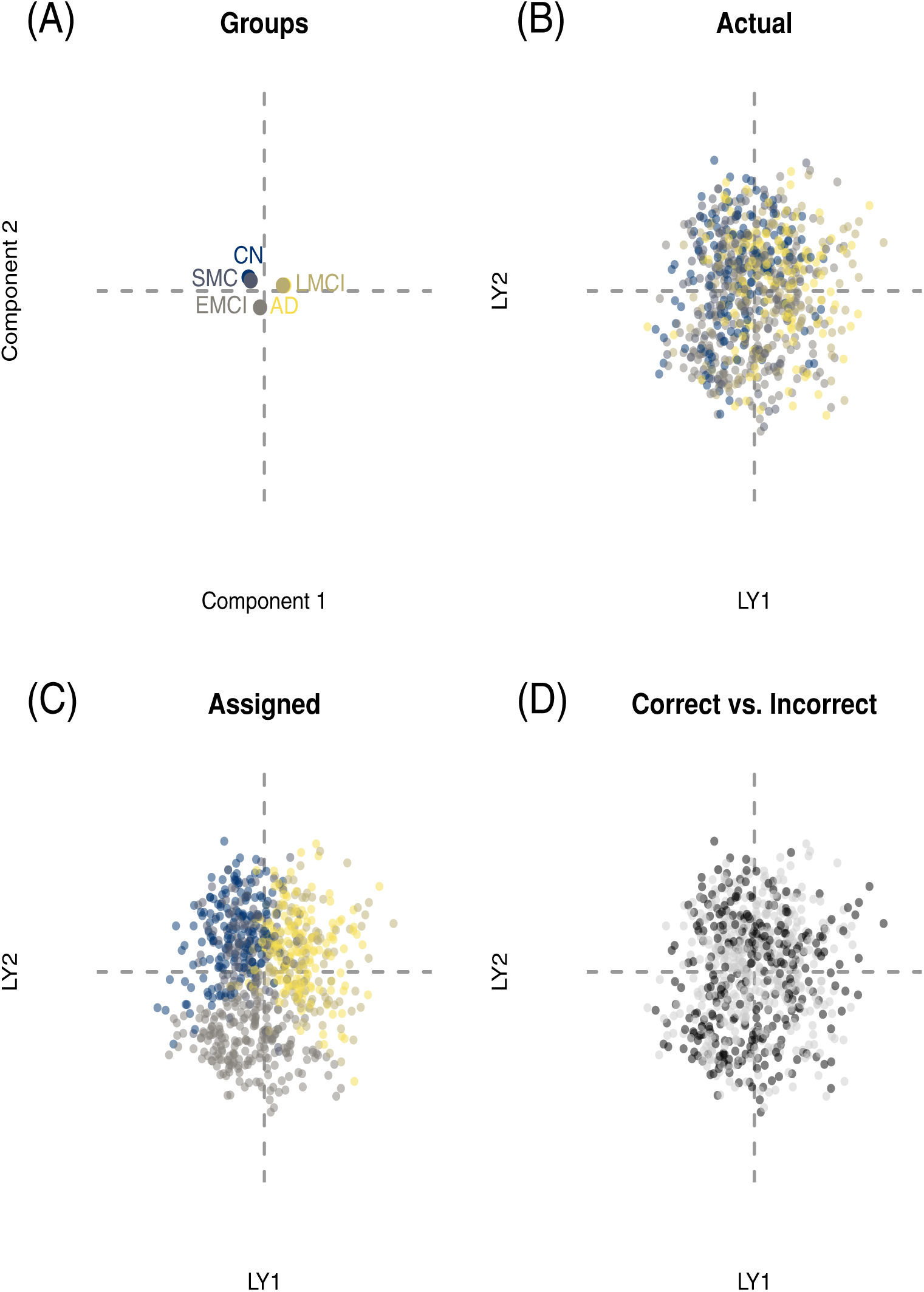
Discriminant PLS-CA-R. (A) shows the component scores for the group on Latent Variables (LV) 1 and 2 (horizontal and vertical respectively), (B) shows the latent variable scores for the genotype (,LY,) LV scores for LVs 1 and 2, colored by *a priori* group association, (C) shows the latent variable scores for the genotype (,LY,) LV scores for LVs 1 and 2, colored by *assigned* group association (i.e., nearest group assignment across all LVs), and (D) shows correct vs. incorrect assignment in black and gray, respectively.

### 3.3 Mixed data and residualization

Our second example illustrates the prediction of genotypes from brain and behavioral variables: (1) three behavioral/clinical scales: Montreal Cognitive Assessment (MoCA) (Nasreddine et al. 2005), Clinical Dementia Rating-Sum of Boxes (CDRSB) (Morris 1993), and Alzheimer’s Disease Assessment Scale (ADAS13) (Skinner et al. 2012), (2) volumetric brain measures in mm^3^: hippocampus (HIPPO), ventricles (VENT), and whole brain (WB), and (3) global estimates of brain function via PET scans: average FDG (for cerebral blood flow; metabolism) in angular, temporal, and posterior cingulate and average AV45 (A*β* tracer) standard uptake value ratio (SUVR) in frontal, anterior cingulate, precuneus, and parietal cortex relative to the cerebellum. This example higlights two features of PLS-CA-R: (1) the ability to accomodate mixed data types (continuous, ordinal, and categorical) and (2) as a way to residualize (orthogonalize; cf. Eq. 17) with respect to known or assumed covariates.

Here, the predictors encompass a variety of data types: all of the brain markers (volumetric MRI estimates, functional PET estimates) and the ADAS13 are quantitative variables, whereas the MoCA and the CDRSB can be considered as ordinal data. Continuous and ordinal data types can be coded into what is called thermometer (Beaton et al. 2018), fuzzy, or “bipolar” coding (because it has two poles) (Greenacre 2014)—an idea initially propsosed by Escofier for continuous data (Escofier 1979). The “Escofier transform” allows continuous data to be analyzed by CA and produces the same results as PCA (Escofier 1979). The same principles can be applied to ordinal data as well (Beaton et al. 2018). Continuous and ordinal data can be transformed into a “pseudo-disjunctive” format that behaves like complete disjunctive data (see Table 1) but preserves the values (as opposed to binning, or dichotomizing). Here, we refer to the transform for continuous data as the “Escofier transform” or “Escofier coding” (Beaton et al. 2016) and the transform for ordinal data as the “thermometer transform” or “thermometer coding”. Because continuous, ordinal, and categorical data can all be transformed into a disjunctive-like format, they can all be analyzed with PLS-CA-R.

The next example identifies the relationship between brain and behavioral markers of AD and genetics. We use the brain and behavioral data as the predictors and the genetics as responses. However, both data sets have their own covariates (some of which are confounds): age, sex, and education influence the behavioral and brain data, where sex, and population origins influence the genotypic variables. For genetic covariates, we use proxies of population origins: self-identified race and ethnicity categories. Both sets of covariates are mixtures of data types (e.g., sex is categorical, age is generally continuous). So, in this example, we illustrate the mixed analysis in two ways—unadjusted and then adjusted for these covariates. First we show the effects of the covariates on the separate data sets, and then compare and contrast adjusted vs. unadjusted analyes. For these analyses, the volumetric brain data were also normalized (divided by) by intracranial volume prior to these analyses to create proportions of total volume for each brain structure.

First we show the PLS-CA-R between each data set and their respective covariates. The main effects of age, sex, and education explained 11.17% of the variance of the behavioral and brain data, where the main effects of sex, race, and ethnicity explained 2.1% of the variance of the genotypic data. The first two components of each analysis are shown in Figure 4. In the brain and behavioral data, age explains a substantial amount of variance for Component 1. In the genotypic analysis, self-identified race and ethnicity are the primary overall effects, where the first two components are explained by those that identify as black or African-American (Component 1) vs. those that identify as Asian, Native, Hawaiian, and/or Latino/Hispanic (Component 2). Both data sets were reconstituted (i.e., **Y**_*ϵ*_ from Eq. 17) from their residuals.

**Figure 4:**
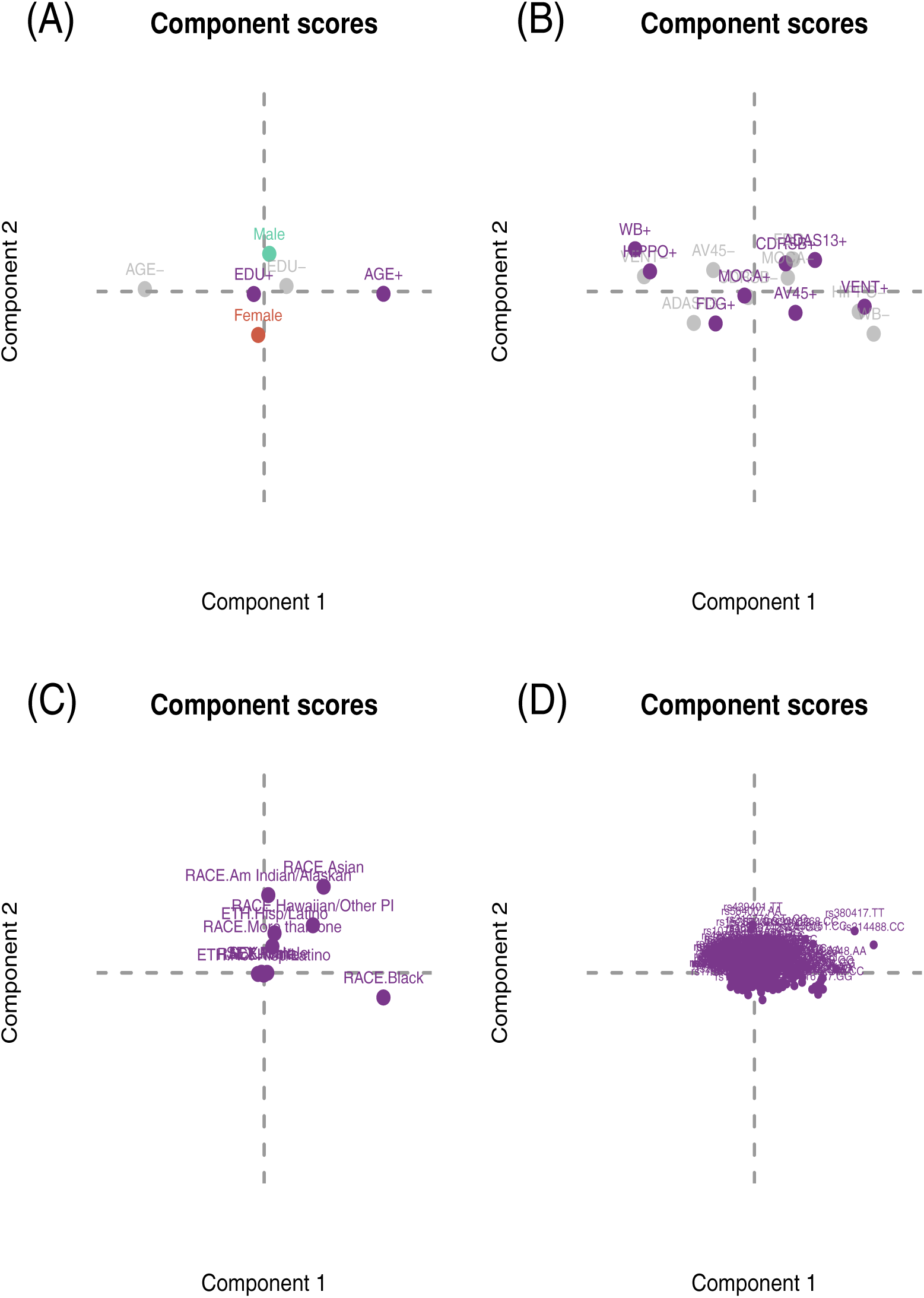
PLS-CA-R used as a way to residualize (orthogonalize) data. The top figures (A) and (B) show prediction of the brain and behavior markers from age, sex, and education. Gray items are one side (lower end) of the “bipolar” or pseudo-disjunctive variables. The bottom figures (C) and (D) show the prediction of genotypes from sex, race, and ethnicity.

Next we performed two analyses with the same goal: to understand the relationship between genetics and behavioral and brain markers of AD. In the unadjusted analysis, the brain and behavioral data explained 1.6% of variance in the genotypic data, whereas in the adjusted analysis, the brain and behavioral data explained 1.54% of variance in the genotypic data. The first two components of the PLS-CA-R results can be seen in Figure 4.

In the unadjusted analysis (Figures 4a and c) vs. the adjusted analysis (Figures 4b and d), we can see some similarities and differences, especially with respect to the behavioral and brain data. AV45 shows little change after residualization, and generally explains a substantial amount of variance as it contributes highly to the first two components in both analyses. The effects of the structural data—especially the hippocampus—are dampened after adjustment (see Figures 4a vs b), where the effects of FDG and CDRSB are now (relatively) increased (see Figures 4a vs b). On the subject level, the differences are small but noticeable, especially for distinguishing between groups (see Figure 6). One important effect is that on a spectrum from CON to AD, we can see that the residualization has a larger impact on the CON side, where the AD side remains somewhat homgeneous (see Figure 6c) for the brain and behavioral variables. With respect to the genotypic LV, there is much less of an effect (see Figure 6d), wherein the observations appear relatively unchanged. However, both pre- (horizontal axis; Figure 6d) and post- (vertical axis; Figure 6d) residualization shows that there are individuals with unique genotypic patterns that remain unaffected by the residualization process (i.e., those at the tails).

**Figure 6:**
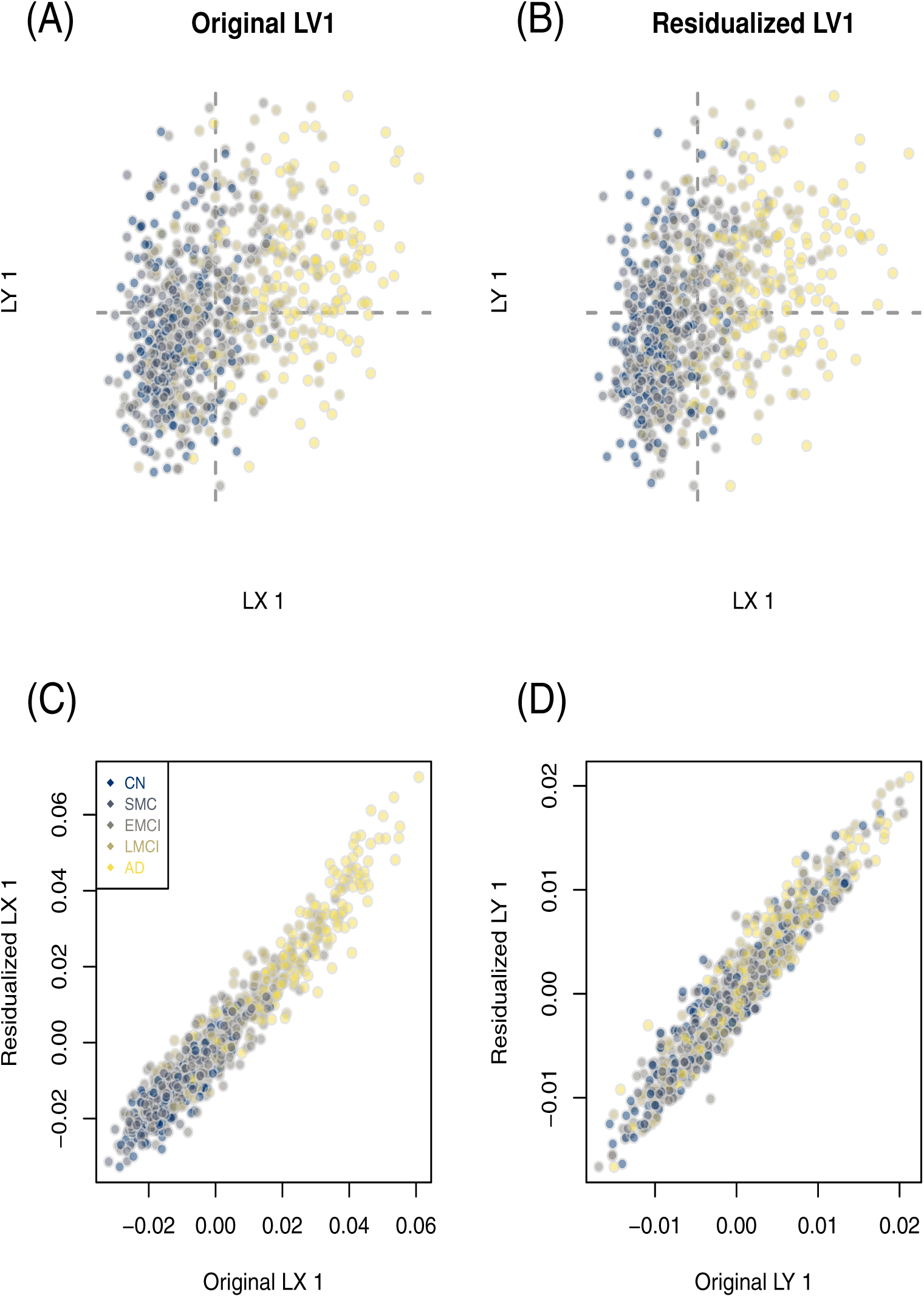
Latent variable scores (observations) for the first latent variable. The top figures (A) and (B) show the projection of the latent variable scores from each set: LX are the brain and behavioral markers, where as LY are the genotypes, for the original and residualized, respectively. The bottom figures (C) and (D) show the the original and residualized scores for the first latent variable compared to one another for each set: the brain and behavioral markers (LX) and the genotypes (LY), respectively.

From this point forward we emphasize the results from the adjusted analyses because they are more realistic in terms of how analyses are performed. For this we refer to Figure 6b—which shows the latent variable scores for the observations and the averages of those scores for the groups—and Figures 5b and 5d—which show, respectively, the component scores for the brain and behavioral markers and the component scores for the genotypes. The first latent variable (Fig. 6b) shows a gradient from control (CON) on the left to Alzheimer’s Disease (AD) on the right. Brain and behavioral variables on the right side of the first component (horizontal axis in Fig. 5b) are more associated with genotypes on the right side (Fig. 5d), where brain and behavioral variables on the left side of the horizontal axis are more associated with genotypes on the left side. In particular, the AA genotype of rs769449, GG genotype of rs2075650, GG genotype of rs4420638, and AA genotype of rs157582 (amongst others) are related to increased AV45 (AV45+), decreased FDG (FDG-), and increased ADAS13 scores (ADAS13+), whereas the TT genotype of rs405697, GG genotype of rs157580, and TC+TT genotypes of rs7412 (amongst others) are more associated with control or possibly protective effects (i.e., decreased AV4, increased FDG, and decreased ADAS13 scores).

**Figure 5:**
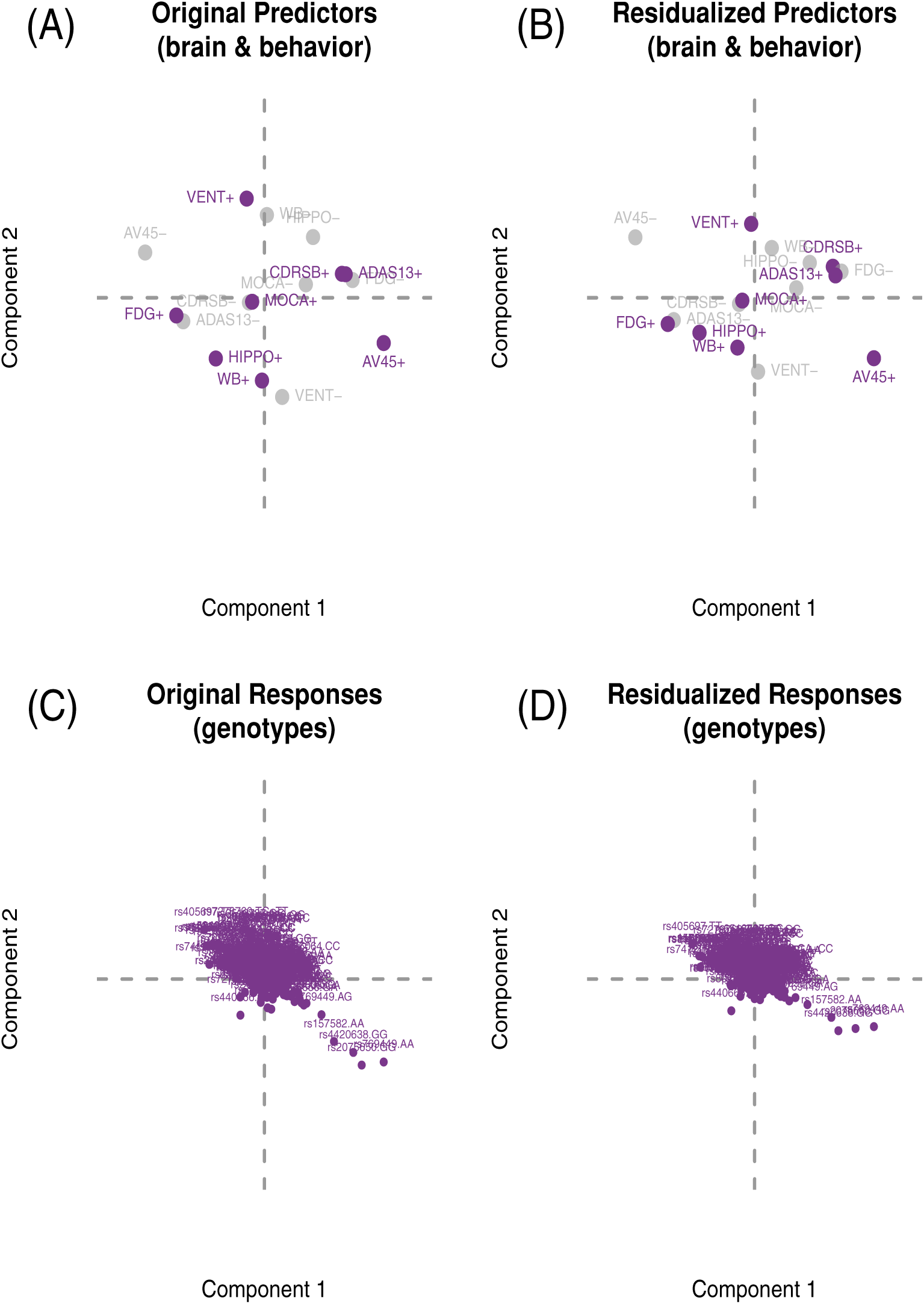
PLS-CA-R to predict genotypes from brain and behavioral markers on the original and residualized data shown on the first two latent variables (components). The top figures (A) and (B) show the component scores for the brain and behavioral markers for the original and residualized data, respectively, and the bottom figures (C) and (D) show the component scores for the genotypes for the original and residualized data, respectively.

### 3.4 SUVR and genotypes

In this final example we use all of the features of PLS-CA-R: mixed data types within and between data sets, each with covariates (and thus require residualization). The goal of this example is to predict genotypes from *β*—amyloid burden (“AV45 uptake”) across regions of the cortex. In this case, because the AV45 uptake data are are strictly non-negative, we effectively treat these data as counts; that is, we would normally apply CA directly to this one data table. Of course, this is only one possible way to handle such data: It is possible to treat these data as row-wise proportions (i.e., percentage of total uptake per region within each subject) or even as continuous data.

Because not all subjects have complete AV45 and genotypic data, the sample for this example is slightly smaller: *N =* 778. Ethnicity, race, and sex (all categorical) explains 2.07% of the variance in the genotypic data where age (numeric), education (ordinal), and sex (categorical) together explain 2.22% of the variance in the in the AV45 uptake data. Overall, AV45 brain data explains 9.08% of the variance in the genotypic data. With the adjusted data we can now perform our intended analyses. Although this analysis produced 67 components (latent variables), we only focus on the first (0.57% of genotypic variance explained by AV45 brain data).

The first latent variable in Figure 7a is only associated with the horizontal axes (Component 1) in Figure 7b and c. The horizontal axis in Fig. 7a is associated with the horizontal axis in Fig. 7b whereas the vertical axis in Fig. 7a is associated with the horizontal axis in Fig. 7c. The first latent variable (Figure 7a) shows a gradient: from left to right we see the groups configured from CN to AD. On the first latent variable we do also see a group-level dissociation where AD+LMCI are entirely on one side whereas EMCI+SMC+CN are on the opposite side for both L_**X**_ (AV45 uptake, horizontal) and **L**_**Y**_ (genotypes, vertical). Higher relative AV45 uptake for the regions on the left side of Component 1 are more associated with EMCI, SMC, and CN than with the other groups, whereas higher relative AV45 uptake for the regions on the right side of Component 1 are more associated with AD and LMCI (Fig. 7b). The genotypes on the left side are associated with the uptake in regions on the left side; likewise the genotypes on the right side are associated with the uptake in regions on the right side (Fig. 7c). For example, LV/Component 1 shows relative uptake in right and left frontal pole, rostral middle frontal, and medial orbitofrontal regions are more associated with the following genotypes: AA and AG from rs769449, GG from rs2075650, GG from rs4420638, and AA from rs157582, than with other genotypes; these effects are generally driven by the AD and LMCI groups. Conversely, LV/Component 1 shows higher relative uptake in right and left lingual, cuneus, as well left parahippocampal and left entorhinal are more associated with the following genotypes: TT from rs405697, GG from rs6859, TC+TT from rs7412, TT from rs2830076, GG from rsl57580, and AA from rs4420638 genotypes than with other genotypes; these effects are generally driven by the GN, SMC, and EMCI cohorts. In summary, the PLS-CA-R results show that particular patterns of regional AV45 uptake predict particular genotypic patterns across many SNPs, and that the sources these effects are generally driven by the groups. Furthermore the underlying brain and genotypic effects of the groups exist along a spectrum of severity.

**Figure 7:**
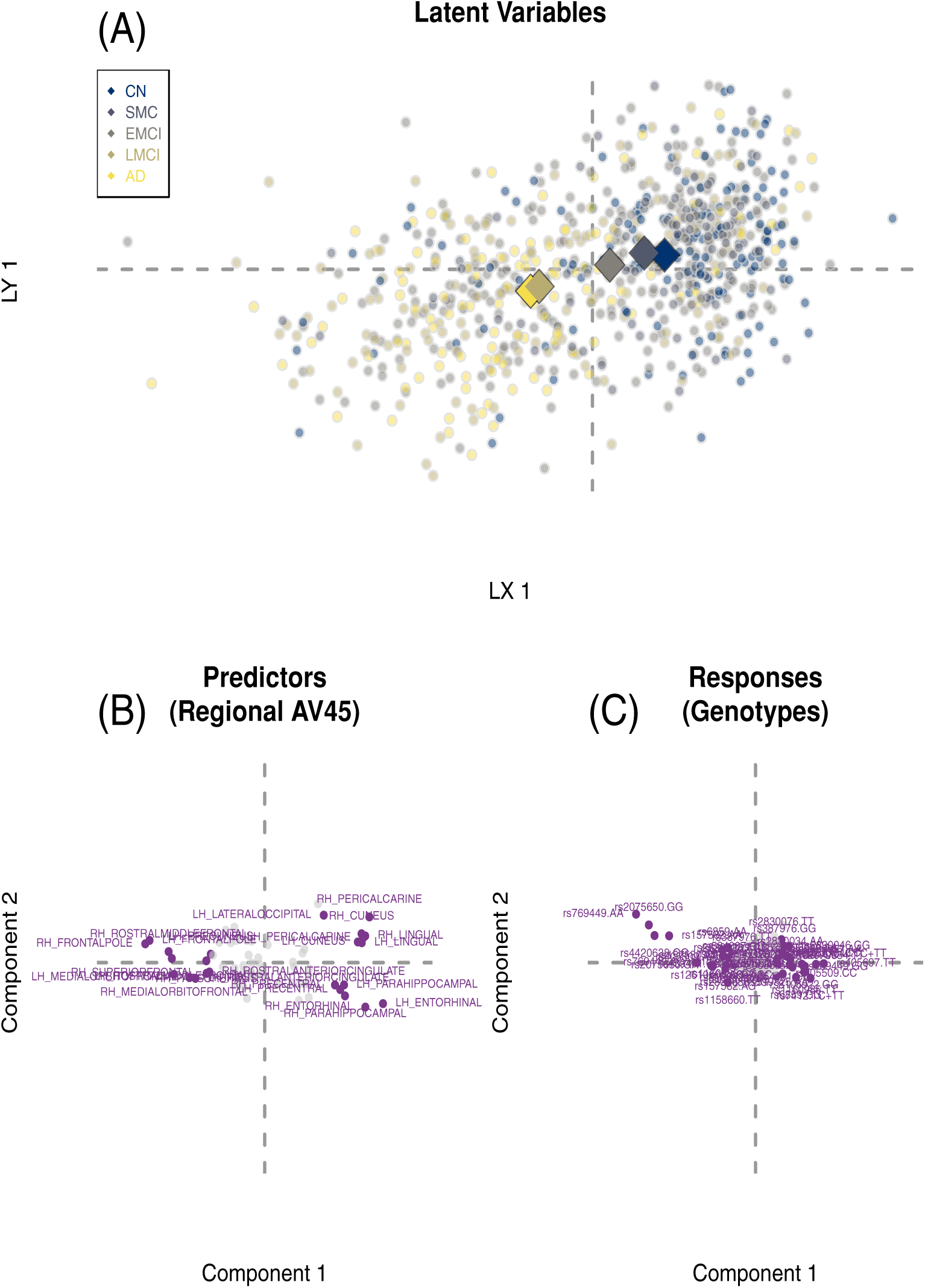
PLS-CA-R to predict genotypes from amyloid burden (“AV45 uptake”). The top figure (A) shows the latent variable scores for the observations on the first latent variable with group averages. The bottom figures (B) and (C) show the amyloid burden in cortical regions and the genotypes, respecively. In (A) we see a gradient from the Alzheimer’s Disease (AD) group to the control (CON) group. Items above expected contribution to variance on the first LV are in purple.

## 4 Discussion

Many modern studies, like ADNI, measure individuals with a variety of scales such as: genetics and genomics, brain structure and function, many aspects of cognition and behavior, batteries of clinical measures, and so on. These data are complex, heterogeneous, and more frequently these data are “wide” (many more variables than subjects) instead of “big” (more subjects than variables). But many current strategies and approaches to handle such multivariate heterogeneous data often requires compromises or sacrifices (e.g., the presumption of single numeric model for categorical data such as the additive model for SNPs; z-scores of ordinal values; or “dichotomania” (https://www.fharrell.eom/post/errmed/#catg): the binning of continuous values into categories). Many of these strategies and approaches presume data are interval scale. Because of the many features and flexibility of PLS-CA-R—e.g., best fit to predictors, orthogonal latent variables, accommodation for virtually any data type—we are able to identify distinct variables and levels (e.g., genotypes) that define or contribute to control (CON) vs. disease (AD) effects (e.g., Fig. 2) or reveal particular patterns anchored by the polar control and disease effects (CON → SMC → EMCI → LMCI → AD; see, e.g., Fig. 7).

While we focused on particular ways of coding and transforming data, there are many alternatives that could be used with PLS-CA-R. For example, we used a disjunctive approach for SNPs because they are categorical—a pattern that matches the genotypic model. However, through various disjunctive schemes, or other forms of Escofier or fuzzy coding, we could have used any genetic model: if all SNPs were coded as the major vs. the minor allele (‘AA’ vs. {‘Aa+aa’}), this would be the dominant model, or we could have assumed the additive model —i.e., 0, 1, 2 for ‘AA’, ‘Aa’, and ‘aa’, respectively—and transformed the data with the ordinal approach (but *not* the continuous approach). We previously provided a comprehensive guide on how to transform various SNP genetic models for use in PLS-CA and CA elsewhere (see Appendix of Beaton et al. 2016). Furthermore, we only highlighted one of many possible methods to transform ordinal data. The term “fuzzy coding” applies more generally to the recoding of ordinal, ranked, preference, or even continuous data. The “fuzzy coding” approach is itself fuzzy, and includes a variety of transformation schemes, all of which conform to the same properties as disjunctive data. The many “fuzzy” and “doubling” coding schemes are described in Escoher (1979), Lebart et al. (1984), or Greenacre (2014). However, for ordinal data—especially with fewer than or equal to 10 levels, and without excessively rare (≤ 1%) Occurences—we recommend to treat ordinal values as categorical levels. When ordinal data are treated as categorical (and disjunctively coded), greater detail about the levels emerges and in most cases could reveal non-linear patterns of the ordinal levels.

We introduced PLS-CA-R in a way that emphasizes various recoding schemes to accomodate different data types. We designed PLS-CA-R as a mixed-data generalization of PLSR, which provides one strategy to identify latent variables and perform regression when standard assumptions are not met (e.g., “high dimensional low sample size” or high collinearity). PLS-CA-R—and GPLS—addresses the need of many fields that require *data type general* methods across multi-source and multi-domain data sets where we require careful considerations about how we prepare and understand our data (Nguyen & Holmes 2019). We introduced PLS-CA-R in a way that emphasizes various recoding schemes to accomodate different data types all with respect to CA and the χ^2^ model. PLS-CA-R provides key features necessary for data analyses as data-rich and data-heavy disciplines and fields rapidly move towards and depend on fundamental techniques in machine and statistical learning (e.g., PLSR, CCA). Finally, with techniques such as mixed-data MFA (Bécue-Bertaut & Pagès 2008), PLS-CA-R provides a much needed basis for development of future methods designed for such complex data sets.

We provide an `R` package with data, documentation, and examples that implements PLS-CA-R (and other cross-decomposition methods) here: https://github.com/derekbeaton/gpls.

Finally, we want to remind the reader to see the Supplemental Material, where we show (1) how PLS-CA-R generalizes most cross-decomposition techniques, (2) alternate PLS algorithms, (3) suggestions on alternate metrics, and finally (4) a ridge-like regularization approach that applies to PLS-CA-R and all the techniques that PLS-CA-R generalizes.

## 4.1 Acknowledgements

Data collection and sharing for this project was funded by the Alzheimer’s Disease Neuroimaging Initiative (ADNI) (National Institutes of Health Grant U01 AG024904) and DOD ADNI (Department of Defense award number W81XWH-12-2-0012). ADNI is funded by the National Institute on Aging, the National Institute of Biomedical Imaging and Bioengineering, and through generous contributions from the following: AbbVie, Alzheimer’s Association; Alzheimer’s Drug Discovery Foundation; Araclon Biotech; BioClinica, Inc.; Biogen; Bristol-Myers Squibb Company; CereSpir, Inc.; Cogstate; Eisai Inc.; Elan Pharmaceuticals, Inc.; Eli Lilly and Company; EuroImmun; F. Hoffmann-La Roche Ltd and its affiliated company Genentech, Inc.; Fujirebio; GE Healthcare; IXICO Ltd.; Janssen Alzheimer Immunotherapy Research & Development, LLC.; Johnson & Johnson Pharmaceutical Research & Development LLC.; Lumosity; Lundbeck; Merck & Co., Inc.; Meso Scale Diagnostics, LLC.; NeuroRx Research; Neurotrack Technologies; Novartis Pharmaceuticals Corporation; Pfizer Inc.; Piramal Imaging; Servier; Takeda Pharmaceutical Company; and Transition Therapeutics. The Canadian Institutes of Health Research is providing funds to support ADNI clinical sites in Canada. Private sector contributions are facilitated by the Foundation for the National Institutes of Health (www.fnih.org). The grantee organization is the Northern California Institute for Research and Education, and the study is coordinated by the Alzheimer’s Therapeutic Research Institute at the University of Southern California. ADNI data are disseminated by the Laboratory for Neuro Imaging at the University of Southern California.

## References

Abdi, H. (2010), ‘Partial least squares regression and projection on latent structure regression (PLS Regression)’, Wiley Interdisciplinary Reviews: Computational Statistics 2(1), 97–106. URL : https://onlinelibraryWileycom/doii/abs/10100wics

Abdi, H. & Béra,. (2018), Correspondence analysis., in R. Alhajj & J. Rokne, eds, ‘Encyclopedia of Social Networks and Mining (2nd Edition)’, Springer Verlag, New York.

Bécue-Bertaut,. & Pagès, J. (2008), ‘Multiple factor analysis and clustering of a mixture of quantitative, categorical and frequency data’, Computational Statistics & Data Analysis 52(6), 32–3268. URL : http://wwwsciencedirectcom/science/article/pii/S016794730700360X

Beaton, D., Dunlop, J., Abdi, H. & Altheimer’s Disease Neuroimaging Initiative (2016), ‘Partial Least Squares Correspondence Analysis: A Framework to Simultaneously Analyte Behavioral and Genetic Data’, Psychological Methods 21(4), 621–6 1.

Beaton, D., Sunderland, K. ., Levine, B., Mandzia, J., Masellis, .M., Swartt, R. H., Troyer, A. K., Binns, . A., Abdi, H., Strother, S. C. & others (2018), ‘Generalitation of the minimum covariance determinant algorithm for categorical and mixed data types’, bioRxiv p. 333005.

Bennet, A.M ., Reynolds, C. A., Gatz, M., Blennow, K., Pedersen, N. L. & Prince, J. A. (2010), ‘Pleiotropy in the presence of allelic heterogeneity: alternative genetic models for the influence of APOE on serum LDL, CSF amyloid-*β*42, and dementia’, Jjournall of Alzheimer ‘s disease: JAD 22(1), 129–134.

Benzécri, J. P. (1973), L’ analyse des doinnées: L’ analyse des correspondances, Dunod.

Bürkner, P.-C. & Vuorre, M. (2019), ‘Ordinal Regression Models in Psychology: A Tutorial’, Advances n Methods and Practices in Psychological Science. Publisher: SAGE PublicationsSage CA: Los Angeles, CA. URL : https://jjournallssagepubcom/doii/101177/2515459188199

Cruchaga, C., Chakraverty, S., Mayo, K., Vallania, F. L. M., Mitra, R. D., Faber, K., Williamson, J., Bird, T., Diaz-Arrastia, R., Foroud, T.M ., Boeve, B. F., Graff-Radford, N. R., St. Jean, P., Lawson, M., Ehm, M. G., Mayeux, R., Goate, A. M. & for the NIA-LOAD/ NCRAD Family Study Consortium (2012), ‘Rare Variants in APP, PSEN1 and PSEN2 Increase Risk for AD in Late-Onset Altheimer’s Disease Families’, PLoS ONE 7(2), e31039. URL : http:dxdoiiorg/10.1371/journalpone003109

Desikan, R. S., Schork, A. J., Wang, Y., Witoelar, A., Sharma, M., McEvoy, L. K., Holland, D., Brewer, J. B., Chen, C.-H., Thompson, W. K., Harold, D., Williams, J., Owen, . J., O’Donovan, M. C., Pericak-Vance, M. A., Mayeux, R., Haines, J. L., Farrer, L. A., Schellenberg, G. D., Heutink, P., Singleton, A. B., Brice, A., Wood, N. W., Hardy, J., Martinez, M., Choi, S. H., DeStefano, A., Ikram, M. A., Bis, J. C., Smith, A., Fitzpatrick, A. L., Launer, L., van Duijn, C., Seshadri, S., Ulstein, I. D., Aarsland, D., Fladby, T., Djurovic, S., Hyman, B. T., Snaedal, J., Stefansson, H., Stefansson, K., Gasser, T., Andreassen, O. A. & Dale, A. M. (2015), ‘Genetic overlap between Alzheimer’s disease and Parkinson’s disease at the MAPT locus’, Molecular Psychiatry. URL : http://wwwnaturecom/mp/journal/vaop/ncurrent/full/mp20156ahtml

Duchesne, S., Chouinard, I., Potvin, O., Fonov, V. S., Khademi, A., Bartha, R., Bellec, P., Collins, D. L., Descoteaux, M., Hoge, R., McCreary, C. R., Ramirez, J., Scott, C. J. M., Smith, E. E., Strother, S. C. & Black, S. E. (2019), ‘The Canadian Dementia Imaging Protocol: Harmoniting National Cohorts’, Journal of Magnetic Resonance imaging 49(2), 456–46. URL : https://onlinebraryWileycom/doi/abs/101002/jmri.26197

Escofier, B. (1979), ‘Traitement simultané de variables qualitatives et quantitatives en analyse factorielle’, Cahiers de Analyse des Doinné es Tome 4(2), 137–146. URL : http://wwwnumdamorg/itemCAD_1979__4_2_137_0/

Escofier, B. (1983), ‘Analyse de la diffé rence entre deux mesures dé finies sur le produit de deux mêmes ensembles’, Cahiers des Analyse des Doinnées Tome 8(3), 325–329. Dunod-Gauthier-Villars. URL : http://wwwnumdamorgitemCAD_1983__8_3_325_0/

Escofier, B. (1984), ‘Analyse factorielle en reférence à un modéle. Application à lanalyse de tableaux dechanges’, Revue de Statistique Appliquée Tome 32(4), 25–-36. Société de statistique de France.URL : http://wwwnumdamorgitemRSA_1984__32_4_25_0/

Escofier-Cordier, B. (1965), L’Analyse des Correspondences., Thése, Faculté des Sciences de Rennes, Université de Rennes. Reprinted in Cahiers du Bureau universitaire de recherche opérationnelle Série Recherche, Tome 13 (1969), pp. 25–59.URL : http://wwwnumdamorgitemBURO_1969__13_25_0/

Farhan, S. M. K., Bartha, R., Black, S. E., Corbett, D., Finger, E., Freedman, M., Greenberg, B., Grimes, D. A., Hegele, R. A., Hudson, C., Kleinstiver, P. W., Lang, A. E., Masellis, M., McIlroy, W. E., McLaughlin, P. M., Montero-Odasso, M., Nunoz, D. G., Munot, D. P., Strother, S., Swartz, R. H., Symons, S., Tartaglia, C., Zinman, L. & Strong, M. J. (2016), ‘The Ontario Neurodegenerative Disease Research Initiative (ONDRI)’, Canadian Journal of Neurological Sciences pp. 1–7. URL: https://www.cambridge.org/core/journals/canadian-journal-of-neurologicalsciences/article/div-classtitlethe-ontario-neurodegenerative-disease-research-initiativeondridiv/3D9558108D69BBF1A4B158DCF2EF6329/core-reader

Genin, E., Hannequin, D., Wallon, D., Sleegers, K., Hiltunen, M., Combarros, O., Bullido, M. J., Engelborghs, S., De Deyn, P., Berr, C., Pasquier, F., Dubois, B., Tognoni, G., Fiévet, N., Brouwers, N., Bettens, K., Arosio, B., Coto, E., Del Zompo, M., Mateo, I., Epelbaum, J., Frank-Garcia, A., Helisalmi, S., Porcellini, E., Pilotto, A., Forti, P., Ferri, R., Scarpini, E., Siciliano, G., Solfrizzi, V., Sorbi, S., Spalletta, G., Valdivieso, F.,Vepsäläinen, S., Alvarez, V., Bosco, P., Mancuso, M., Panza, F., Nacmias, B., Bossù, P., Hanon, O., Piccardi, P., Annoni, G., Seripa, D., Galimberti, D., Licastro, F., Soininen, H., Dartigues, J.-F., Kamboh, M. I., Van Broeckhoven, C., Lambert, J. C., Amouyel, P. & Campion, D. (2011), ‘APOE and Alzheimer disease: a major gene with semi-dominant inheritance’, Molecular psychiatry 16(9), 903–907.

Grassi, M. & Visentin, S. (1994), ‘Correspondence analysis applied to grouped cohort data’, Statistics in Medicine 13(23-24), 2407–2425.

Greenacre, M. (2014), Data Doubling and Fuzzy Coding, in J. Blasius & M. Greenacre, eds, ‘Visualization and Verbalization of Data’, CRC Press, Philadelphia, PA, USA, pp. 239–253.

Greenacre, M. J. (1984), Theory and Applications of Correspondence Analysis, Academic Press.

Greenacre, M. J. (1993), ‘Biplots in correspondence analysis’, Journal of Applied Statistics 20(2), 251–269.

Greenacre, M. J. (2010), ‘Correspondence analysis’, Wiley Interdisciplinary Reviews: Computational Statistics 2(5), 613–619. URL: https://onlinelibrary.wiley.com/doi/abs/10.1002/wics.114

Huang, Y.-W. A., Zhou, B., Wernig, M. & Südhof, T. C. (2017), ‘ApoE2, ApoE3, and ApoE4 Di_erentially Stimulate APP Transcription and Aβ Secretion’, Cell. URL://www.sciencedirect.com/science/article/pii/S0092867416317603

Jonsson, T., Atwal, J. K., Steinberg, S., Snaedal, J., Jonsson, P. V., Bjornsson, S., Stefansson, H., Sulem, P., Gudbjartsson, D., Maloney, J., Hoyte, K., Gustafson, A., Liu, Y., Lu, Y., Bhangale, T., Graham, R. R., Huttenlocher, J., Bjornsdottir, G., Andreassen, O. A., Jönsson, E. G., Palotie, A., Behrens, T. W., Magnusson, O. T., Kong, A., Thorsteinsdottir, U., Watts, R. J. & Stefansson, K. (2012), ‘A mutation in APP protects against Alzheimer’s disease and age-related cognitive decline’, Nature 488(7409), 96–99. URL: https://www.nature.com/articles/nature11283

Lebart, L., Morineau, A. & Warwick, K. M. (1984), Multivariate descriptive statistical analysis: correspondence analysis and related techniques for large matrices, Wiley.

Lettre, G., Lange, C. & Hirschhorn, J. N. (2007), ‘Genetic model testing and statistical power in population-based association studies of quantitative traits’, Genetic Epidemiology: The Offcial Publication of the International Genetic Epidemiology Society 31(4), 358–362.

Linnertz, C., Anderson, L., Gottschalk, W., Crenshaw, D., Lutz, M. W., Allen, J., Saith, S., Mihovilovic, M., Burke, J. R., Welsh-Bohmer, K. A., Roses, A. D. & Chiba-Falek, O. (2014), ‘The cis-regulatory effect of an Alzheimer’s disease-associated poly-T locus on expression of TOMM40 and apolipoprotein E genes’, Alzheimer’s & Dementia 10(5), 541–551. URL: http://www.sciencedirect.com/science/article/pii/S155252601302801X

Montero-Odasso, M., Pieruccini-Faria, F., Bartha, R., Black, S. E., Finger, E., Freedman, M., Greenberg, B., Grimes, D. A., Hegele, R. A., Hudson, C., Kleinstiver, P. W., Lang, A. E., Masellis, M., McLaughlin, P. M., Munoz, D. P., Strother, S., Swartz, R. H., Symons, S., Tartaglia, M. C., Zinman, L., Strong, M. J., ONDRI Investigators & McIlroy, W. (2017), ‘Motor Phenotype in Neurodegenerative Disorders: Gait and Balance Platform Study Design Protocol for the Ontario Neurodegenerative Research Initiative (ONDRI)’, Journal of Alzheimer’s disease: JAD 59(2), 707–721.

Morris, J. C. (1993), ‘The Clinical Dementia Rating (CDR): current version and scoring rules.’, Neurology 43(11), 2412–2414. Lippincott Williams & Wilkins.

Myers, A. J., Kaleem, M., Marlowe, L., Pittman, A. M., Lees, A. J., Fung, H. C., Duckworth, J., Leung, D., Gibson, A., Morris, C. M., Silva, R. d. & Hardy, J. (2005), ‘The H1c haplotype at the MAPT locus is associated with Alzheimer’s disease’, Human Molecular Genetics 14(16), 2399–2404. URL: http://hmg.oxfordjournals.org/content/14/16/2399

Nasreddine, Z. S., Phillips, N. A., Bédirian, V., Charbonneau, S., Whitehead, V., Collin, I., Cummings, J. L. & Chertkow, H. (2005), ‘The Montreal Cognitive Assessment, MoCA: A Brief Screening Tool For Mild Cognitive Impairment’, Journal of the American Geriatrics Society 53(4), 695–699. URL: https://onlinelibrary.wiley.com/doi/abs/10.1111/j.1532-5415.2005.53221.x

Nguyen, L. H. & Holmes, S. (2019), ‘Ten quick tips for effective dimensionality reduction’, PLOS Computational Biology 15(6), e1006907.

Peterson, D., Munger, C., Crowley, J., Corcoran, C., Cruchaga, C., Goate, A. M., Norton, M. C., Green, R. C., Munger, R. G., Breitner, J. C. S., Welsh-Bohmer, K. A., Lyketsos, C., Tschanz, J. & Kauwe, J. S. K. (2014), ‘Variants in PPP3r1 and MAPT are associated with more rapid functional decline in Alzheimer’s disease: The Cache County Dementia Progression Study’, Alzheimer’s & Dementia 10(3), 366–371. URL: http://www.sciencedirect.com/science/article/pii/S1552526013000861

Pérez-Enciso, M. & Tenenhaus, M. (2003), ‘Prediction of clinical outcome with microarray data: a partial least squares discriminant analysis (PLS-DA) approach’, Human Genetics 112(5), 581–592. URL: https://doi.org/10.1007/s00439-003-0921-9

Purcell, S., Neale, B., Todd-Brown, K., Thomas, L., Ferreira, M. A., Bender, D., Maller, J., Sklar, P., De Bakker, P. I., Daly, M. J. et al. (2007), ‘Plink: a tool set for wholegenome association and population-based linkage analyses’, The American journal of human genetics 81(3), 559–575.

Rodríguez-Pérez, R., Fernández, L. & Marco, S. (2018), ‘Overoptimism in cross-validation when using partial least squares-discriminant analysis for omics data: a systematic study’, Analytical and Bioanalytical Chemistry 410(23), 5981–5992. URL: https://doi.org/10.1007/s00216-018-1217-1

Roses, A. D., Lutz, M. W., Amrine-Madsen, H., Saunders, A. M., Crenshaw, D. G., Sundseth, S. S., Huentelman, M. J., Welsh-Bohmer, K. A. & Reiman, E. M. (2010), ‘A TOMM40 variable-length polymorphism predicts the age of late-onset Alzheimer’s disease’, The Pharmacogenomics Journal 10(5), 375–384.

Skinner, J., Carvalho, J. O., Potter, G. G., Thames, A., Zelinski, E., Crane, P. K. & Gibbons, L. E. (2012), ‘The Alzheimer’s Disease Assessment Scale-Cognitive-Plus (ADASCog-Plus): an expansion of the ADAS-Cog to improve responsiveness in MCI’, Brain imaging and behavior 6(4). URL: https://www.ncbi.nlm.nih.gov/pmc/articles/PMC3873823/

Tenenhaus, M. (1998), La régression PLS: théorie et pratique, Editions TECHNIP.

Trabzuni, D., Wray, S., Vandrovcova, J., Ramasamy, A., Walker, R., Smith, C., Luk, C., Gibbs, J. R., Dillman, A., Hernandez, D. G., Arepalli, S., Singleton, A. B., Cookson, M. R., Pittman, A. M., Silva, R. d., Weale, M. E., Hardy, J. & Ryten, M. (2012), ‘MAPT expression and splicing is differentially regulated by brain region: relation to genotype and implication for tauopathies’, Human Molecular Genetics 21(18), 4094–4103. URL: http://hmg.oxfordjournals.org/content/21/18/4094

Vormfelde, S. V. & Brockmöller, J. (2007), ‘On the value of haplotype-based genotype _phenotype analysis and on data transformation in pharmacogenetics and -genomics’, Nature Reviews Genetics 8(12). URL: http://www.nature.com.libproxy.utdallas.edu/nrg/journal/v8/n12/full/nrg1916-c1.html

Wold, H. (1975), ‘Soft modelling by latent variables: the non-linear iterative partial least squares (NIPALS) approach’, Journal of Applied Probability 12(S1), 117–142. Cambridge University Press.

Wold, S., Esbensen, K. & Geladi, P. (1987), ‘Principal component analysis’, Chemometrics and Intelligent Laboratory Systems 2(1), 37–52. URL: http://www.sciencedirect.com/science/article/pii/0169743987800849

Wold, S., Ruhe, A., Wold, H. & Dunn, III, W. (1984), ‘The collinearity problem in linear regression. The partial least squares (PLS) approach to generalized inverses’, SIAM Journal on Scientific and Statistical Computing 5(3), 735–743.

Wold, S., Sjöström, M. & Eriksson, L. (2001), ‘PLS-regression: a basic tool of chemometrics’, Chemometrics and Intelligent Laboratory Systems 58(2), 109–130. URL: http://www.sciencedirect.com/science/article/pii/S0169743901001551

