## Supplemental Material for "A generalization of partial least squares regression and correspondence analysis for categorical and mixed data: An application with the ADNI data"

July 16, 2020

**Abstract**

We provide supplemental material for partial least squares-correspondence analysis-regression (PLS-CA-R_ that highlights how PLS-CA-R provides the baiss for generalization of numerous cross-decomposition methods (e.g., PLS, CCA, RRR) as well as ridge-like regularization. Here we provide additional material and explanations (e.g., algorithms) based on the formulation in the main text.

*Keywords:* generalized partial least squares, canonical correlation analysis, reduced rank regression, ridge regularization, R package

*Data used in preparation of this article were obtained from the Alzheimer's Disease Neuroimaging Initiative (ADNI) database (`http://adni.loni.usc.edu/`). As such, the investigators within the ADNI contributed to the design and implementation of ADNI and/or provided data but did not participate in analysis or writing of this report. A complete listing of ADNI investigators can be found at http://adni.loni.ucla.edu/wpcontent/uploads/how_to_apply/ADNI_Acknowledgement_List.pdf

### 1 Introduction

This Appendix provides additional details on PLS-CA-R and, in particular, a variety of extensions of and generalizations from PLS-CA-R. We also provide additional details on some concepts, such as the generalized singular value decomposition. As established in the main text, PLS-CA-R provides a generalization of PLS-R for categorical and mixed data. However, PLS-CA-R provides the basis for futher generalizations that extend to other optimizations, alternate metrics, different PLS algorithms, and even ridge-like regularization. In this Appendix we explain those additional generalizations and variations based on how we established PLS-CA-R in the main text.

First, we explain the relationship between the SVD and GSVD in a more detail than in the main text. Following that, we extend the concept of the GSVD triplet for PLS with what we call "the GPLSSVD sextuplet". Second, we show how the GPLSSVD triplet allows us to perform PLS-CA-R, as well as other cross-decomposition techniques, more easily. From there, we use the GPLSSVD triplet as a way to further simplify the three primary PLS algorithms (regression, correlation, canonical). Third we provide a short discussion on how the GPLSSVD—which is inspired from PLS-CA-R—gives us a more unified way to accomodate different optimizations (e.g., partial least squares vs. canonical correlation) and different weights or metrics. We then present two ways to perform a ridge-like regularization with an emphasis on PLS-CA-R, but the regularization applies to any technique under the GPLSSVD framework. Finally, we point out numerous strategies and approaches for inference and stability assessment that are easily adapted for PLS-CA-R specifically and GPLSSVD generally.

### 2 The SVD, GSVD, and GPLSSVD

The SVD of $\mathbf{X}$ is

$$\mathbf{X} = \mathbf{U}\boldsymbol{\Delta}\mathbf{V}^T, \tag{1}$$

where $\mathbf{U}^T\mathbf{U} = \mathbf{I} = \mathbf{V}^T\mathbf{V}$. The GSVD of $\mathbf{X}$ is

$$\mathbf{X} = \mathbf{P}\boldsymbol{\Delta}\mathbf{Q}^T, \tag{2}$$

where $\mathbf{P}^T\mathbf{M}\mathbf{P} = \mathbf{I} = \mathbf{Q}^T\mathbf{W}\mathbf{Q}$. Practically, the GSVD is performed through the SVD as $\widetilde{\mathbf{X}} = \mathbf{M}^{\frac{1}{2}}\mathbf{X}\mathbf{W}^{\frac{1}{2}} = \mathbf{U}\boldsymbol{\Delta}\mathbf{V}^T$, where the generalized singular vectors are computed from the singular vectors as $\mathbf{P} = \mathbf{M}^{-\frac{1}{2}}\mathbf{U}$ and $\mathbf{Q} = \mathbf{W}^{-\frac{1}{2}}\mathbf{V}$. The relationship between the SVD and GSVD can be expressed through what we decompose as

$$\widetilde{\mathbf{X}} = \mathbf{M}^{\frac{1}{2}}\mathbf{X}\mathbf{W}^{\frac{1}{2}} \Longleftrightarrow \mathbf{X} = \mathbf{M}^{-\frac{1}{2}}\widetilde{\mathbf{X}}\mathbf{W}^{-\frac{1}{2}}. \quad (3)$$

As noted in the main text, the GSVD can be presented in "triplet notation" as GSVD($\mathbf{M}, \mathbf{X}, \mathbf{W}$).

#### 2.1 From GSVD triplet to GPLSSVD sextuplet

We introduce an extension of the GSVD triplet for PLS, called the "GPLSSVD sextuplet". The GPLSSVD sextuplet helps us in two ways: (1) it simplifies some of the notation and decomposition concepts, and (2) it provides the basis for generalization of cross-decomposition methods (e.g., canonical correlation, reduced rank regression).

The "GPLSSVD sextuplet" takes the form of GPLSSVD($\mathbf{M}_{\mathbf{X}}, \mathbf{Z}_{\mathbf{X}}, \mathbf{W}_{\mathbf{X}}, \mathbf{M}_{\mathbf{Y}}, \mathbf{Z}_{\mathbf{Y}}, \mathbf{W}_{\mathbf{Y}}$) and like the GSVD, decomposes a matrix $\mathbf{Z}_{\mathbf{R}} = (\mathbf{M}_{\mathbf{X}}^{\frac{1}{2}}\mathbf{Z}_{\mathbf{X}})^T\mathbf{M}_{\mathbf{Y}}^{\frac{1}{2}}\mathbf{Z}_{\mathbf{Y}}$ as

$$\mathbf{Z}_{\mathbf{R}} = \mathbf{P}\boldsymbol{\Delta}\mathbf{Q}^T \text{ with } \mathbf{P}^T\mathbf{W}_{\mathbf{X}}\mathbf{P} = \mathbf{I} = \mathbf{Q}^T\mathbf{W}_{\mathbf{Y}}\mathbf{Q}. \quad (4)$$

From this decomposition, the GPLSSVD produces latent variables as

$$\mathbf{L}_{\mathbf{X}} = \mathbf{Z}_{\mathbf{X}}\mathbf{W}_{\mathbf{X}}\mathbf{P} \text{ and } \mathbf{L}_{\mathbf{Y}} = \mathbf{Z}_{\mathbf{Y}}\mathbf{W}_{\mathbf{Y}}\mathbf{Q} \text{ where } \mathbf{L}_{\mathbf{X}}^T\mathbf{L}_{\mathbf{Y}} = \boldsymbol{\Delta}. \quad (5)$$

Alternatively, we can show the GPLSSVD through the SVD. Let us refer to $\widetilde{\mathbf{Z}}_{\mathbf{X}} = \mathbf{M}_{\mathbf{X}}^{\frac{1}{2}}\mathbf{Z}_{\mathbf{X}}\mathbf{W}_{\mathbf{X}}^{\frac{1}{2}}$ and $\widetilde{\mathbf{Z}}_{\mathbf{Y}} = \mathbf{M}_{\mathbf{Y}}^{\frac{1}{2}}\mathbf{Z}_{\mathbf{Y}}\mathbf{W}_{\mathbf{Y}}^{\frac{1}{2}}$, where $\widetilde{\mathbf{Z}}_{\mathbf{R}} = \widetilde{\mathbf{Z}}_{\mathbf{X}}^T\widetilde{\mathbf{Z}}_{\mathbf{Y}} = (\mathbf{M}_{\mathbf{X}}^{\frac{1}{2}}\mathbf{Z}_{\mathbf{X}}\mathbf{W}_{\mathbf{X}}^{\frac{1}{2}})^T(\mathbf{M}_{\mathbf{Y}}^{\frac{1}{2}}\mathbf{Z}_{\mathbf{Y}}\mathbf{W}_{\mathbf{Y}}^{\frac{1}{2}})$. We decompose $\widetilde{\mathbf{Z}}_{\mathbf{R}}$ as

$$\widetilde{\mathbf{Z}}_{\mathbf{R}} = \mathbf{U}\boldsymbol{\Delta}\mathbf{V}^T \text{ with } \mathbf{U}^T\mathbf{U} = \mathbf{I} = \mathbf{V}^T\mathbf{V}. \quad (6)$$

We can also compute the latent variables as

$$\mathbf{L}_{\mathbf{X}} = \widetilde{\mathbf{Z}}_{\mathbf{X}}\mathbf{U} \text{ and } \mathbf{L}_{\mathbf{Y}} = \widetilde{\mathbf{Z}}_{\mathbf{Y}}\mathbf{V} \text{ where } \mathbf{L}_{\mathbf{X}}^T\mathbf{L}_{\mathbf{Y}} = \boldsymbol{\Delta}. \quad (7)$$

Like with the SVD and GSVD, the GPLSSVD has the same orthogonality constraints: $\mathbf{U}^T\mathbf{U} = \mathbf{I} = \mathbf{V}^T\mathbf{V}$, or $\mathbf{P}^T\mathbf{M}_{\mathbf{Y}}\mathbf{P} = \mathbf{I} = \mathbf{Q}^T\mathbf{W}_{\mathbf{Y}}\mathbf{Q}$. The relationship between the singular vectors are $\mathbf{U} = \mathbf{W}_{\mathbf{X}}^{\frac{1}{2}}\mathbf{P} \iff \mathbf{P} = \mathbf{M}_{\mathbf{X}}^{-\frac{1}{2}}\mathbf{U}$ and $\mathbf{V} = \mathbf{W}_{\mathbf{Y}}^{\frac{1}{2}}\mathbf{Q} \iff \mathbf{Q} = \mathbf{W}_{\mathbf{Y}}^{-\frac{1}{2}}\mathbf{V}$. We can also compute component (a.k.a. factor) scores from the GPLSSVD as $\mathbf{F}_J = \mathbf{W}_{\mathbf{X}}\mathbf{P}\boldsymbol{\Delta}$ and $\mathbf{F}_K = \mathbf{W}_{\mathbf{Y}}\mathbf{Q}\boldsymbol{\Delta}$. An alternate form of the component scores are $\mathbf{F}'_J = \mathbf{W}_{\mathbf{X}}\mathbf{P}$ and $\mathbf{F}'_K = \mathbf{W}_{\mathbf{Y}}\mathbf{Q}$.

Finally, we note one last extension to the GPLSSVD to make it the "GPLSSVD septuplet" (similarly we also a "GSVD quadruplet"). We can include desired rank to return as an input parameter for the GPLSSVD septuplet (and GSVD quadruplet) as GPLSSVD($\mathbf{M}_{\mathbf{X}}, \mathbf{Z}_{\mathbf{X}}, \mathbf{W}_{\mathbf{X}}, \mathbf{M}_{\mathbf{Y}}, \mathbf{Z}_{\mathbf{Y}}$[illegible] (and GSVD($\mathbf{M}_{\mathbf{X}}, \mathbf{Z}_{\mathbf{X}}, \mathbf{W}_{\mathbf{X}}, C$)), where $C$ is an integer to indicate the rank of the solution desired. For example, when $C = 1$ the GPLSSVD (and GSVD) return a rank 1 solution. The $C$ parameter could be any value (constrained to the minimum rank of either $\mathbf{X}$ or $\mathbf{Y}$), but $C = 1$ is particlarly convenient for our following generalizations.

### 3 PLS and GPLS algorithms

Though we have presented PLS-CA regression as a generalization of PLS regression that accomodates virutally any data type (by way of CA), the way we formalized PLS-CA regression leads to further variants and broader generalizations. These generalizations span (1) various PLS, CA, and related approaches, (2) several typical PLS algorithms, (3) a variety of optimizations (e.g., canonical correlation), and (4) ridge-like regularization.

There exist three commonly used PLS algorithms: (1) PLS regression (PLS-REG) decomposition (Wold 1975, Wold et al. 1984, 2001, Abdi 2010), (2) PLS correlation (PLS-COR) decomposition (Bookstein 1994, Ketterlinus et al. 1989) generally more known in neuroimaging (McIntosh et al. 1996, McIntosh & Lobaugh 2004, Krishnan et al. 2011) and also has numerous alternate names such as PLS-SVD, co-inertia (Dolédec & Chessel 1994, Dray (2014)), and Tucker's interbattery factor analysis (Tucker 1958) amongst others (see also Beaton et al. 2016), and (3) PLS canonical (PLS-CAN) decomposition (Tenenhaus 1998, Wegelin et al. 2000) which is a symmetric method (like PLS-COR) with iterative deflation (like PLS-REG).

Based on how we formalized PLS-CA regression, we now show how PLS-CA regression provides the basis of generalizations of these three algorithms, as well as further optimizations, similar to Borga et al. (1992), Indahl et al. (2009), and de Micheaux et al. (2019). But we do so in a more comprehensive way that incorporates more methods than other unification strategies, and we also do so in a way that accomodates multiple data types. We refer to the generalization of the three previously mentioned PLS techniques under the umbrella of generalized partial least squares (GPLS) as GPLS-COR, GPLS-REG, and GPLS-CAN, for the "correlation", "regression", and "canonical" decompositions respectively. GPLS-COR and GPLS-CAN are symmetric decomposition approaches where neither $\mathbf{X}$ nor $\mathbf{Y}$ are privileged. GPLS-REG is an asymmetric decomposition approach where $\mathbf{X}$ is privileged. We present the GPLS-COR, GPLS-REG, and then GPLS-CAN algorithms with their respective optimizations. We do so in the previously mentioned order because GPLS-COR—by way of the GPLSSVD—is used as the basis of all three algorithms and GPLS-CAN shares features and concepts with both GPLS-COR and GPLS-REG. For all of these we rely on the formlization of PLS-CA regression as established in the main text.

#### 3.1 GPLS-COR

The GPLS-COR decomposition is the simplest GPLS technique. It requires only a single pass of the SVD—or in our case the GPLSSVD. There are no explicit iterative steps in GPLS-COR. GPLS-COR takes as input the two preprocessed matrices—$\mathbf{Z_X}$ and $\mathbf{Z_Y}$—and their respective row and column weights: $\mathbf{M_X}$ and $\mathbf{W_X}$ for $\mathbf{Z_X}$, and $\mathbf{M_Y}$ and $\mathbf{W_Y}$ for $\mathbf{Z_Y}$, where $C$ is the desired number of components to return. GPLS-COR is shown in Algorithm 1.

GPLS-COR maximizes the relationship between $\mathbf{L_X}$ and $\mathbf{L_Y}$ with the orthogonality constraint $\boldsymbol{\ell}_{\mathbf{X},c}^T \boldsymbol{\ell}_{\mathbf{Y},c'} = 0$ when $c \neq c'$ where $\boldsymbol{\ell}_{\mathbf{X},c}^T \boldsymbol{\ell}_{\mathbf{Y},c} = \delta_c$ and thus $\mathbf{L}_{\mathbf{X}}^T \mathbf{L_Y} = \mathbf{U}^T \widetilde{\mathbf{Z}}_{\mathbf{X}}^T \widetilde{\mathbf{Z}}_{\mathbf{Y}} \mathbf{V}^T = \mathbf{U}^T \widetilde{\mathbf{Z}}_{\mathbf{R}} \mathbf{V}^T = \mathbf{U}^T \mathbf{U} \boldsymbol{\Delta} \mathbf{V}^T \mathbf{V}^T = \boldsymbol{\Delta}$. We can also show this with the generalized vectors and constraints as $\mathbf{L}_{\mathbf{X}}^T \mathbf{L_Y} = \mathbf{P}^T \mathbf{W_X} \mathbf{Z}_{\mathbf{X}}^T \mathbf{M}_{\mathbf{X}}^{\frac{1}{2}} \mathbf{M}_{\mathbf{Y}}^{\frac{1}{2}} \mathbf{Z_Y} \mathbf{W_Y} \mathbf{Q}^T = \mathbf{P}^T \mathbf{W_X} \mathbf{P} \boldsymbol{\Delta} \mathbf{Q}^T \mathbf{W_Y} \mathbf{Q} = \boldsymbol{\Delta}$.

The primary example of GPLS-COR is the standard "PLS correlation" approach. Let's assume that $\mathbf{X}$ and $\mathbf{Y}$ are comprised of continuous data. So, $\mathbf{Z_X}$ and $\mathbf{Z_Y}$ are column-wise centered (and/or normalized) versions of $\mathbf{X}$ and $\mathbf{Y}$. When all weight matrices are identity matrices, then the GPLSSVD implements the "PLS correlation" decomposition as GPLSSVD($\mathbf{I}, \mathbf{Z_X}, \mathbf{I}, \mathbf{I}, \mathbf{Z_Y}, \mathbf{I}$) and also as shown in Algorithm 1. However, the way we estab-

**Result:** Generalized PLS-correlation between $\mathbf{Z_X}$ and $\mathbf{Z_Y}$

**Input :** $\mathbf{M_X}$, $\mathbf{Z_X}$, $\mathbf{W_X}$, $\mathbf{M_Y}$, $\mathbf{Z_Y}$, $\mathbf{W_Y}$, $C$

**Output:** $\mathbf{\Delta}$, $\mathbf{U}$, $\mathbf{V}$, $\mathbf{P}$, $\mathbf{Q}$, $\mathbf{F}_J$, $\mathbf{F}_K$, $\mathbf{L_X}$, $\mathbf{L_Y}$

GPLSSVD($\mathbf{M_X}, \mathbf{Z_X}, \mathbf{W_X}, \mathbf{M_Y}, \mathbf{Z_Y}, \mathbf{W_Y}, C$)

**Algorithm 1:** Generalized PLS-correlation algorithm. GPLS-COR is the GPLSSVD and provides the basis of other GPLS techniques. Furthermore, GPLS-COR easily allows for a variety of optmizations for examples canonical correlation, reduced rank regression (redundancy analysis), and even ridge-like regularization, which then extend to the other GPLS algorithms (i.e., regression and canonical decompositions). Note that this is a truncated version of the algorithm and does not include all of the GPLSSVD outputs.

| Method | $\mathbf{M_X}$ | $\mathbf{W_X}$ | $\mathbf{M_Y}$ | $\mathbf{W_Y}$ |
|---|---|---|---|---|
| PLS-COR | $\mathbf{I}$ | $\mathbf{I}$ | $\mathbf{I}$ | $\mathbf{I}$ |
| RRR/RDA | $\mathbf{I}$ | $(\mathbf{Z_X}^T\mathbf{Z_X})^{-1}$ | $\mathbf{I}$ | $\mathbf{I}$ |
| CCA | $\mathbf{I}$ | $(\mathbf{Z_X}^T\mathbf{Z_X})^{-1}$ | $\mathbf{I}$ | $(\mathbf{Z_Y}^T\mathbf{Z_Y})^{-1}$ |

Table 1: The primary cross-decomposition techniques framed as GPLSSVD approaches.

lished the GPLSSVD and Algorithm 1 allows us to obtain the results of three of the most common cross-decomposition ("two-table") techniques: PLS correlation (shown above), reduced rank regression (RRR, a.k.a., reduced rank regression [RDA]), and canonical correlation analysis (CCA). RRR/RDA is performed as GPLSSVD($\mathbf{I}, \mathbf{Z_X}, (\mathbf{Z_X}^T\mathbf{Z_X})^{-1}, \mathbf{I}, \mathbf{Z_Y}, \mathbf{I}$) and CCA is performed as GPLSSVD($\mathbf{I}, \mathbf{Z_X}, (\mathbf{Z_X}^T\mathbf{Z_X})^{-1}, \mathbf{I}, \mathbf{Z_Y}, (\mathbf{Z_Y}^T\mathbf{Z_Y})^{-1}$). In Table 1, we show how these three techniques are related through the GPLSSVD by highlighting that they are merely a change in choice of weights (metrics).

Furthermore, these three variants—PLSC, CCA, and RDA/RRR—also generalize discriminant analyses under different optimizations so long either $\mathbf{X}$ or $\mathbf{Y}$ (depending on the technique) is a dummy-coded (complete disjunctive) matrix, where each observation (row) is assigned to a specific group or category (columns). Going further, if we have disjunctive (or pseudo-disjunctive) data—as we do in the main text— we can use GPLSSVD to obtain the results of PLS-CA "correlation" (Beaton et al. 2016). Using the same matrices as established in the main text, PLS-CA "correlation" is GPLSSVD($\mathbf{M}_{\mathbf{X}}^{-1}, \mathbf{Z}_{\mathbf{X}}, \mathbf{W}_{\mathbf{X}}^{-1}, \mathbf{M}_{\mathbf{Y}}^{-1}, \mathbf{Z}_{\mathbf{Y}}, \mathbf{W}_{\mathbf{Y}}^{-1}$), where $\mathbf{M}_{\mathbf{X}}$ and $\mathbf{M}_{\mathbf{Y}}$ are diagonal matrices of row frequencies for each matrix, where $\mathbf{W}_{\mathbf{X}}$ and $\mathbf{W}_{\mathbf{Y}}$ are diagonal matrices of column frequencies for each matrix, and $\mathbf{Z}_{\mathbf{X}}$ and $\mathbf{Z}_{\mathbf{Y}}$ are the deviations from independence matrices.

GPLS-COR—which is just the GPLSSVD—provides the basis for the other two algorithms: both GPLS-REG and GPLS-CAN make use of GPLS-COR (i.e., the GPLSSVD) with rank 1 solutions iteratively.

#### 3.2 GPLS-REG

The GPLS-REG decomposition uses the GPLSSVD septuplet iteratively for $C$ iterations, with only a rank 1 solution is provided for each use of the GPLSSVD. Alternatively, GPLS-REG can be thought of a direct extension of GPLS-COR as defined in Algorithm 1.

Then the two data matrices—$\mathbf{Z}_{\mathbf{X}}$ and $\mathbf{Z}_{\mathbf{Y}}$—are deflated for each step asymmetrically, with a privileged $\mathbf{Z}_{\mathbf{X}}$. GPLS-REG is shown in Algorithm 2.

**Result:** Generalized PLS-regression between $\mathbf{Z}_{\mathbf{X}}$ and $\mathbf{Z}_{\mathbf{Y}}$
**Input :** $\mathbf{M}_{\mathbf{X}}$, $\mathbf{Z}_{\mathbf{X}}$, $\mathbf{W}_{\mathbf{X}}$, $\mathbf{M}_{\mathbf{Y}}$, $\mathbf{Z}_{\mathbf{Y}}$, $\mathbf{W}_{\mathbf{Y}}$, $C$
**Output:** $\widetilde{\mathbf{\Delta}}$, $\widetilde{\mathbf{U}}$, $\widetilde{\mathbf{V}}$, $\widetilde{\mathbf{P}}$, $\widetilde{\mathbf{Q}}$, $\widetilde{\mathbf{F}}_J$, $\widetilde{\mathbf{F}}_K$, $\mathbf{L}_{\mathbf{X}}$, $\mathbf{L}_{\mathbf{Y}}$, $\mathbf{T}_{\mathbf{X}}$, $\widehat{\mathbf{U}}$, $\mathbf{B}$

**for** $c = 1, \ldots, C$ **do**
  GPLSSVD($\mathbf{M}_{\mathbf{X}}, \mathbf{Z}_{\mathbf{X}}, \mathbf{W}_{\mathbf{X}}, \mathbf{M}_{\mathbf{Y}}, \mathbf{Z}_{\mathbf{Y}}, \mathbf{W}_{\mathbf{Y}}, 1$)
  $\mathbf{t}_{\mathbf{X}} \leftarrow \boldsymbol{\ell}_{\mathbf{X}} \times ||\boldsymbol{\ell}_{\mathbf{X}}||^{-1}$
  $b \leftarrow \boldsymbol{\ell}_{\mathbf{Y}}^T \mathbf{t}_{\mathbf{X}}$
  $\widehat{\mathbf{u}} \leftarrow (\mathbf{M}_{\mathbf{X}}^{\frac{1}{2}} \mathbf{Z}_{\mathbf{X}} \mathbf{W}_{\mathbf{X}}^{\frac{1}{2}})^T \mathbf{t}_{\mathbf{X}}$
  $\mathbf{Z}_{\mathbf{X}} \leftarrow \mathbf{Z}_{\mathbf{X}} - [\mathbf{M}_{\mathbf{X}}^{-\frac{1}{2}} (\mathbf{t}_{\mathbf{X}} \widehat{\mathbf{u}}^T) \mathbf{W}_{\mathbf{X}}^{-\frac{1}{2}}]$
  $\mathbf{Z}_{\mathbf{Y}} \leftarrow \mathbf{Z}_{\mathbf{Y}} - [\mathbf{M}_{\mathbf{Y}}^{-\frac{1}{2}} (b \mathbf{t}_{\mathbf{X}} \mathbf{v}^T) \mathbf{W}_{\mathbf{Y}}^{-\frac{1}{2}}]$
**end**

**Algorithm 2:** Generalized PLS-regression algorithm. The results of a rank 1 GPLSSVD are used to compute the latent variables and values necessary for deflation of $\mathbf{Z}_{\mathbf{X}}$ and $\mathbf{Z}_{\mathbf{Y}}$. Note that this is a truncated version of the algorithm and does not include all of the GPLSSVD outputs.

GPLS-REG maximizes the relationship between $\mathbf{L}_{\mathbf{X}}$ and $\mathbf{L}_{\mathbf{Y}}$ with the orthogonality constraint $\boldsymbol{\ell}_{\mathbf{X},c}^{T}\boldsymbol{\ell}_{\mathbf{X},c'} = 0$ when $c \neq c'$ where $\boldsymbol{\ell}_{\mathbf{X},c}^{T}\boldsymbol{\ell}_{\mathbf{Y},c} = \delta_{c}$ which is also $\text{diag}\{\mathbf{L}_{\mathbf{X}}^{T}\mathbf{L}_{\mathbf{Y}}\} = \text{diag}\{\widetilde{\boldsymbol{\Delta}}\}$. When $\mathbf{Z}_{\mathbf{X}}$ and $\mathbf{Z}_{\mathbf{Y}}$ are column-wise centered (and/or normalized) versions of $\mathbf{X}$ and $\mathbf{Y}$, and when all weight matrices are identity matrices, then GPLS-REG is (one of the) traditional "PLS regression" decomposition(s; akin to the SIMPLS algorithm in Tenenhaus (1998)). We detail the regression decomposition for categorical and mixed data in the main text.

#### 3.3 GPLS-CAN

The GPLS-CAN decomposition can be thought of as a compromise between GPLS-COR and GPLS-REG, it: (1) is symmetric like GPLS-COR, and (2) uses the GPLSSVD septuplet iteratively for $C$ iterations—with only a rank 1 solution is provided for each use of the GPLSSVD—like GPLS-REG. In GPLS-CAN, the two data matrices—$\mathbf{Z}_{\mathbf{X}}$ and $\mathbf{Z}_{\mathbf{Y}}$—are deflated for each iteration. GPLS-CAN is shown in Algorithm 3

GPLS-CAN maximizes the relationship between $\mathbf{L}_{\mathbf{X}}$ and $\mathbf{L}_{\mathbf{Y}}$ with the orthogonality constraints $\boldsymbol{\ell}_{\mathbf{X},c}^{T}\boldsymbol{\ell}_{\mathbf{X},c'} = 0$ and $\boldsymbol{\ell}_{\mathbf{Y},c}^{T}\boldsymbol{\ell}_{\mathbf{Y},c'} = 0$ when $c \neq c'$ where $\boldsymbol{\ell}_{\mathbf{X},c}^{T}\boldsymbol{\ell}_{\mathbf{Y},c} = \delta_{c}$ which is also $\text{diag}\{\mathbf{L}_{\mathbf{X}}^{T}\mathbf{L}_{\mathbf{Y}}\} = \text{diag}\{\widetilde{\boldsymbol{\Delta}}\}$.

#### 3.4 GPLS algorithms summary

Note that across the three algorithms defined here, that the first component is identical when the same preprocessed data and weights (a.k.a. metrics) are provided to the GPLSSVD. In many cases, subsequent components across the three algorithms differ, but generally do not differ substantially. The similarities are because of the common approach to maximization: $\boldsymbol{\ell}_{\mathbf{X},c}^{T}\boldsymbol{\ell}_{\mathbf{Y},c} = \delta_{c}$. The differences are because of the different orthogonality constraints when $c \neq c'$ where: (1) GPLS-COR in Algorithm 1 is $\boldsymbol{\ell}_{\mathbf{X},c}^{T}\boldsymbol{\ell}_{\mathbf{Y},c'} = 0$, (2) GPLS-REG in Algorithm 2 is $\boldsymbol{\ell}_{\mathbf{X},c}^{T}\boldsymbol{\ell}_{\mathbf{X},c'} = 0$, and (3) GPLS-CAN in Algorithm 3 is both $\boldsymbol{\ell}_{\mathbf{X},c}^{T}\boldsymbol{\ell}_{\mathbf{X},c'} = 0$ and $\boldsymbol{\ell}_{\mathbf{Y},c}^{T}\boldsymbol{\ell}_{\mathbf{Y},c'} = 0$.

**Result:** Generalized PLS-canonical between $\mathbf{Z_X}$ and $\mathbf{Z_Y}$

**Input :** $\mathbf{M_X}$, $\mathbf{Z_X}$, $\mathbf{W_X}$, $\mathbf{M_Y}$, $\mathbf{Z_Y}$, $\mathbf{W_Y}$, $C$

**Output:** $\widetilde{\mathbf{U}}$, $\widetilde{\mathbf{V}}$, $\widetilde{\mathbf{P}}$, $\widetilde{\mathbf{Q}}$, $\widetilde{\mathbf{F}}_J$, $\widetilde{\mathbf{F}}_K$, $\mathbf{L_X}$, $\mathbf{L_Y}$, $\widetilde{\mathbf{\Delta}}$, $\mathbf{T_X}$, $\mathbf{T_Y}$, $\widehat{\mathbf{U}}$, $\widehat{\mathbf{V}}$

**for** $c = 1, \ldots, C$ **do**

GPLSSVD$(\mathbf{M_X}, \mathbf{Z_X}, \mathbf{W_X}, \mathbf{M_Y}, \mathbf{Z_Y}, \mathbf{W_Y}, 1)$

$\mathbf{t_X} \leftarrow \boldsymbol{\ell}_\mathbf{X} \times ||\boldsymbol{\ell}_\mathbf{X}||^{-1}$

$\mathbf{t_Y} \leftarrow \boldsymbol{\ell}_\mathbf{Y} \times ||\boldsymbol{\ell}_\mathbf{Y}||^{-1}$

$\widehat{\mathbf{u}} \leftarrow (\mathbf{M}_\mathbf{X}^{\frac{1}{2}} \mathbf{Z_X} \mathbf{W}_\mathbf{X}^{\frac{1}{2}})^T \mathbf{t_X}$

$\widehat{\mathbf{v}} \leftarrow (\mathbf{M}_\mathbf{Y}^{\frac{1}{2}} \mathbf{Z_Y} \mathbf{W}_\mathbf{Y}^{\frac{1}{2}})^T \mathbf{t_Y}$

$\mathbf{Z_X} \leftarrow \mathbf{Z_X} - [\mathbf{M}_\mathbf{X}^{-\frac{1}{2}} (\mathbf{t_X} \widehat{\mathbf{u}}^T) \mathbf{W}_\mathbf{X}^{-\frac{1}{2}}]$

$\mathbf{Z_Y} \leftarrow \mathbf{Z_Y} - [\mathbf{M}_\mathbf{Y}^{-\frac{1}{2}} (\mathbf{t_Y} \widehat{\mathbf{v}}^T) \mathbf{W}_\mathbf{Y}^{-\frac{1}{2}}]$

**end**

**Algorithm 3:** Generalized PLS-canonical algorithm. The results of a rank 1 GPLSSVD are used to compute the latent variables and values necessary for deflation of $\mathbf{Z_X}$ and $\mathbf{Z_Y}$. The deflation in GPLS-CAN is the same as in GPLS-REG in Algorithm 2, but for both matrices. Note that this is a truncated version of the algorithm and does not include all of the GPLSSVD outputs.

#### 3.5 GPLS optimizations and further generalizations

From the GPLS perspective, we can better unify the wide variety of approaches with similar goals but variations of metrics, transformations, and optimizations that often appear under a wide variety of names (e.g., PLS, CCA, interbattery factor analysis, co-inertia analysis, canonical variates, PLS-CA, and so on; see Abdi et al. (2017)). The way we defined the GPLS algorithms—in particular with the weights applied to the rows and columns of each data matrix—leads to numerous further generalizations.

Given the way we establish the GPLS algorithms here, we provide the basis for further generalizations of many approaches. This is especially true for the numerous variants of correspondecne analysis, such as power transformations for CA (Greenacre 2009) alternate distance metrics such as Hellinger distances (Rao 1995, Escofier 1978), or "non-symmetrical CA" (D'Ambra & Lauro 1992, Kroonenberg & Lombardo 1999, Takane et al. 1991). For many of these approaches, the weight matrices—either for the rows or columns of a given matrix—are what change between techniques. In some of the aforementioned cases (e.g., power transformations) there are also additional steps to preprocess the data.

#### 3.6 Ridge-like regularizations

We show two possible strategies for ridge-like regularization for the GPLSSVD (which then applies to any of the algorithms we outline above). We first show these two regularization approaches specifically for the PLS-CA framework. From there we briefly discuss how these regularization approaches extend to other techniques (e.g., CCA, PLS-COR) under the GPLSSVD framework.

The first approach is based on Takane's regularized multiple CA (Takane & Hwang 2006) and regularized nonsymmetric CA (Takane & Jung 2009). To do so, it is convenient to slightly reformulate PLS-CA-R, but we still require $\mathbf{X}$, $\mathbf{Y}$, $\mathbf{O_X}$, $\mathbf{O_Y}$, $\mathbf{E_X}$, and $\mathbf{E_Y}$ as defined in the main text. First we re-define $\mathbf{Z_X} = (\mathbf{O_X} - \mathbf{E_X}) \times (\mathbf{1}^T\mathbf{X}1)$ and $\mathbf{Z_Y} = (\mathbf{O_Y} - \mathbf{E_Y}) \times (\mathbf{1}^T\mathbf{Y}1)$. Next we define the following additional matrices: $\mathbf{D}_{\mathbf{X},I} = \text{diag}\{\mathbf{X1}\}$, and $\mathbf{D}_{\mathbf{Y},I} = \text{diag}\{\mathbf{Y1}\}$ which are diagonal matrices of the row sums of $\mathbf{X}$ and $\mathbf{Y}$, and $\mathbf{D}_{\mathbf{X},J} = \text{diag}\{\mathbf{1}^{\text{T}}\mathbf{X}\}$, and $\mathbf{D}_{\mathbf{Y},K} = \text{diag}\{\mathbf{1}^{\text{T}}\mathbf{Y}\}$ which are the column sums of $\mathbf{X}$ and $\mathbf{Y}$. Then PLS-CA correlation, regression, and canonical decompositions replace the

GPLSSVD step in Algorithms 1, 2, 3 with GPLSSVD($\mathbf{D}_{\mathbf{X},I}^{-1}, \mathbf{Z}_{\mathbf{X}}^{T}, \mathbf{D}_{\mathbf{X},J}^{-1}, \mathbf{D}_{\mathbf{Y},I}^{-1}, \mathbf{Z}_{\mathbf{Y}}^{T}, \mathbf{D}_{\mathbf{Y},K}^{-1}$). The only differences between this reformulation and what we originally established is that the generalized singular vectors ($\mathbf{P}$ and $\mathbf{Q}$) and the component scores ($\mathbf{F}_{\mathbf{J}}$ and $\mathbf{F}_{\mathbf{K}}$) differ by constant scaling factors (which are the sums of $\mathbf{X}$ and $\mathbf{Y}$ for their respective scores).

We can regularize PLS-CA-R in the same way as Takane's RMCA. We require (1) a ridge parameter which we refer to as $\epsilon$ and (2) variants of $\mathbf{D}_{\mathbf{X},I}$, $\mathbf{D}_{\mathbf{X},J}$, $\mathbf{D}_{\mathbf{Y},I}$, and $\mathbf{D}_{\mathbf{Y},K}$ that we refer to as $\mathbb{D}_{\mathbf{X},I} = \mathbf{D}_{\mathbf{X},I} + [\epsilon \times (\mathbf{Z}_{\mathbf{X}}\mathbf{Z}_{\mathbf{X}}^{T})^{+}]$, $\mathbb{D}_{\mathbf{Y},I} = \mathbf{D}_{\mathbf{Y},I} + [\epsilon \times (\mathbf{Z}_{\mathbf{Y}}\mathbf{Z}_{\mathbf{Y}}^{T})^{+}]$, $\mathbb{D}_{\mathbf{X},J} = \mathbf{D}_{\mathbf{X},J} + [\epsilon \times \mathbf{Z}_{\mathbf{X}}^{T}(\mathbf{Z}_{\mathbf{X}}\mathbf{Z}_{\mathbf{X}}^{T})^{+}\mathbf{Z}_{\mathbf{X}}]$, and $\mathbb{D}_{\mathbf{Y},K} = \mathbf{D}_{\mathbf{Y},K} + [\epsilon \times \mathbf{Z}_{\mathbf{Y}}^{T}(\mathbf{Z}_{\mathbf{Y}}\mathbf{Z}_{\mathbf{Y}}^{T})^{+}\mathbf{Z}_{\mathbf{Y}}]$. When $\epsilon = 0$ then $\mathbb{D}_{\mathbf{X},I} = \mathbf{D}_{\mathbf{X},I}$, $\mathbb{D}_{\mathbf{Y},I} = \mathbf{D}_{\mathbf{Y},I}$, $\mathbb{D}_{\mathbf{X},J} = \mathbf{D}_{\mathbf{X},J}$, $\mathbb{D}_{\mathbf{Y},K} = \mathbf{D}_{\mathbf{Y},K}$. We obtain ridge-like regularized forms of PLS-CA for the correlation, regression, and canonical decompositions if we replace the GPLSSVD step in (or just the input to) each algorithm with GPLSSVD($\mathbb{D}_{\mathbf{X},I}^{-1}, \mathbf{Z}_{\mathbf{X}}^{T}, \mathbb{D}_{\mathbf{X},J}^{-1}, \mathbb{D}_{\mathbf{Y},I}^{-1}, \mathbf{Z}_{\mathbf{Y}}^{T}, \mathbb{D}_{\mathbf{Y},K}^{-1}$). As per Takane's recommendation (Takane & Hwang 2006), $\epsilon$ could be any positive value, though integers in the range from 1 to 20 provide sufficient regularization.

However, the above approach may not be feasible when $I$, $J$, and/or $K$ are particularly large because the various crossproduct and projection matrices require a large amount of memory and/or computational expense. So,we can use a "truncated" version of the Takane regularization which is more computationally efficient, and analogous to the regularization procedure of Allen (Allen 2013, Allen et al. 2014). We re-define $\mathbb{D}_{\mathbf{X},I} = \mathbf{D}_{\mathbf{X},I} + (\epsilon \times \mathbf{I})$ and $\mathbb{D}_{\mathbf{Y},I} = \mathbf{D}_{\mathbf{Y},I} + (\epsilon \times \mathbf{I})$ and then also $\mathbb{D}_{\mathbf{X},J} = \mathbf{D}_{\mathbf{X},J} + (\epsilon \times \mathbf{I})$ and $\mathbb{D}_{\mathbf{Y},K} = \mathbf{D}_{\mathbf{Y},K} + (\epsilon \times \mathbf{I})$ where $\mathbf{I}$ are identity matrices of appropriate size. Like in the previous formulation, we replace the values we have in the GPLSSVD step where GPLSSVD($\mathbb{D}_{\mathbf{X},I}^{-1}, \mathbf{Z}_{\mathbf{X}}^{T}, \mathbb{D}_{\mathbf{X},J}^{-1}, \mathbb{D}_{\mathbf{Y},I}^{-1}, \mathbf{Z}_{\mathbf{Y}}^{T}, \mathbb{D}_{\mathbf{Y},K}^{-1}$); and in this particular case, the weight matrices are all diagonal matrices, which allows for a lower memory footprint and less computational burden.

We have two concluding remarks on the ridge-like regularizations we presented. First, though the above are presented under teh PLS-CA frameworks, the basis of these concepts extend to any of the GPLSSVD techniques. In particular, the more simplified Takane/Allen hybrid approach to ridge-like regularization is easier to generally apply: it requires only some inflation factor (i.e., $\epsilon$) along the diagonals of the weight matrices. However, the first (Takane's) approach was established in a framework more akin to CCA, and thus could also be used for any of the optimization approaches outlined in Table 1. Second, though we presented ridge-like regularization with a single $\epsilon$ it is entirely possible to use different $\epsilon$s for each set of weights. Although it is possible, we do not necessarily recommend this approach, as it requires a (potentially expensive) grid search over all the various $\epsilon$ parameters. Alternatively, if multiple $\epsilon$s were used, one could minimize the number of parameters to search and set some of the $\epsilon$s to 0 and, for example, use only one or two $\epsilon$ values instead of four possible $\epsilon$ values.

##### 3.7 Implementation of algorithms

We provide an `R` package that implements all of the algorithms, with the cross-decomposition variations here: `https://github.com/derekbeaton/gpls`. This package provides direct interfaces to PLS-COR, PLS-REG, PLS-CAN, CCA, RRR, and PLS-CA-COR, PLS-CA-REG, and PLS-CA-CAN, as well as the generalized PLS approaches as outlined above.

#### 4 Further extensions of PLS-CA-R and GPLSSVD

In both the main text and here, we have foregone any discussions of inference, stability, and resampling for PLS-CA-R (or GPLSSVD) in part because many of the inference and stability approaches established throughout the broader PLS (and CCA) literature still apply with little or no changes. Such approaches include feature selection or sparsification (Sutton et al. 2018), additional regularization or sparsification approaches (Le Floch et al. 2012, Guillemot et al. 2019, Tenenhaus et al. 2014, Tenenhaus & Tenenhaus 2011), cross-validation (Wold et al. 1987, Rodríguez-Pérez et al. 2018, Kvalheim et al. 2019, Abdi 2010), permutation (Berry et al. 2011, Winkler et al. 2020), various bootstrap approaches (Abdi 2010, Takane & Jung 2009) or tests (McIntosh & Lobaugh 2004, Krishnan et al. 2011), and other frameworks such as split-half resampling (Strother et al. 2002, Kovacevic et al. 2013, Strother et al. 2004).

### References


Abdi, H. (2010), 'Partial least squares regression and projection on latent structure regression (PLS Regression)', *Wiley Interdisciplinary Reviews: Computational Statistics* **2**(1), 97–106.
**URL:** *https://onlinelibrary.wiley.com/doi/abs/10.1002/wics.51*

Abdi, H., Guillemot, V., Eslami, A. & Beaton, D. (2017), 'Canonical correlation analysis', *Encyclopedia of Social Network Analysis and Mining* pp. 1–16. Springer.

Allen, G. I. (2013), 'Sparse and Functional Principal Components Analysis', *arXiv:1309.2895 [stat]* . arXiv: 1309.2895.
**URL:** *http://arxiv.org/abs/1309.2895*

Allen, G. I., Grosenick, L. & Taylor, J. (2014), 'A Generalized Least-Square Matrix Decomposition', *Journal of the American Statistical Association* **109**(505), 145–159.
**URL:** *http://dx.doi.org/10.1080/01621459.2013.852978*

Beaton, D., Dunlop, J., Abdi, H. & Alzheimer's Disease Neuroimaging Initiative (2016), 'Partial Least Squares Correspondence Analysis: A Framework to Simultaneously Analyze Behavioral and Genetic Data', *Psychological Methods* **21**(4), 621–651.

Berry, K. J., Johnston, J. E. & Mielke, P. W. (2011), 'Permutation methods', *Wiley Interdisciplinary Reviews: Computational Statistics* **3**, 527–542.
**URL:** *http://wires.wiley.com/WileyCDA/WiresArticle/wisId-WICS177.html*

Bookstein, F. L. (1994), 'Partial least squares: A dose-response model for measurement in the behavioral and brain sciences.', *Psycoloquy* .

Borga, M., Landelius, T. & Knutsson, H. (1992), *A Unified Approach to PCA, PLS, MLR and CCA*.

D'Ambra, L. & Lauro, N. C. (1992), 'Non symmetrical exploratory data analysis', *Statistica Applicata* **4**(4), 511–529.

de Micheaux, P. L., Liquet, B., Sutton, M. et al. (2019), 'Pls for big data: A unified parallel algorithm for regularised group pls', *Statistics Surveys* **13**, 119–149.

Dolédec, S. & Chessel, D. (1994), 'Co-inertia analysis: an alternative method for studying species-environment relationships', *Freshwater Biology* **31**, 177–194.

Dray, S. (2014), Analysing a pair of tables: Coinertia analysis and duality diagrams, *in* J. Blasius & G. M, eds, 'Visualization and Verbalization of Data', CRC Press, Boca Raton, pp. 289–300.

Escofier, B. (1978), 'Analyse factorielle et distances répondant au principe d'équivalence distributionnelle', *Revue de statistique appliquée* **26**(4), 29–37.

Greenacre, M. (2009), 'Power transformations in correspondence analysis', *Computational Statistics & Data Analysis* **53**(8), 3107–3116.

Guillemot, V., Beaton, D., Gloaguen, A., Löfstedt, T., Levine, B., Raymond, N., Tenenhaus, A. & Abdi, H. (2019), 'A constrained singular value decomposition method that integrates sparsity and orthogonality', *PloS one* **14**(3), e0211463.

Indahl, U. G., Liland, K. H. & Næs, T. (2009), 'Canonical partial least squares—a unified pls approach to classification and regression problems', *Journal of Chemometrics: A Journal of the Chemometrics Society* **23**(9), 495–504.

Ketterlinus, R. D., Bookstein, F. L., Sampson, P. D. & Lamb, M. E. (1989), 'Partial least squares analysis in developmental psychopathology', *Development and Psychopathology* **1**(4), 351–371.

Kovacevic, N., Abdi, H., Beaton, D. & McIntosh, A. R. (2013), Revisiting pls resampling: comparing significance versus reliability across range of simulations, *in* 'New Perspectives in Partial Least Squares and Related Methods', Springer, pp. 159–170.

Krishnan, A., Williams, L. J., McIntosh, A. R. & Abdi, H. (2011), 'Partial Least Squares (PLS) methods for neuroimaging: A tutorial and review', *NeuroImage* **56**(2), 455 – 475. Multivariate Decoding and Brain Reading.
**URL:** *http://www.sciencedirect.com/science/article/pii/S1053811910010074*

Kroonenberg, P. M. & Lombardo, R. (1999), ‘Nonsymmetric correspondence analysis: A tool for analysing contingency tableswith a dependence structure’, *Multivariate Behavioral Research* **34**(3), 367–396.

Kvalheim, O. M., Grung, B. & Rajalahti, T. (2019), ‘Number of components and prediction error in partial least squares regression determined by Monte Carlo resampling strategies’, *Chemometrics and Intelligent Laboratory Systems* .
**URL:** *http://www.sciencedirect.com/science/article/pii/S0169743918307056*

Le Floch, E., Guillemot, V., Frouin, V., Pinel, P., Lalanne, C., Trinchera, L., Tenenhaus, A., Moreno, A., Zilbovicius, M., Bourgeron, T., Dehaene, S., Thirion, B., Poline, J.-B. & Duchesnay, E. (2012), ‘Significant correlation between a set of genetic polymorphisms and a functional brain network revealed by feature selection and sparse Partial Least Squares’, *NeuroImage* **63**(1), 11–24.
**URL:** *http://www.sciencedirect.com/science/article/pii/S1053811912006775*

McIntosh, A., Bookstein, F., Haxby, J. & Grady, C. (1996), ‘Spatial Pattern Analysis of Functional Brain Images Using Partial Least Squares’, *NeuroImage* **3**(3), 143–157.
**URL:** *http://www.sciencedirect.com/science/article/pii/S1053811996900166*

McIntosh, A. R. & Lobaugh, N. J. (2004), ‘Partial least squares analysis of neuroimaging data: applications and advances’, *Neuroimage* **23**, S250–S263.

Rao, C. R. (1995), ‘A review of canonical coordinates and an alternative to correspondence analysis using Hellinger distance’, *Qüestiió: quaderns d’estadística i investigació operativa* **19**(1).

Rodríguez-Pérez, R., Fernández, L. & Marco, S. (2018), ‘Overoptimism in cross-validation when using partial least squares-discriminant analysis for omics data: a systematic study’, *Analytical and Bioanalytical Chemistry* **410**(23), 5981–5992.
**URL:** *https://doi.org/10.1007/s00216-018-1217-1*

Strother, S. C., Anderson, J., Hansen, L. K., Kjems, U., Kustra, R., Sidtis, J., Frutiger, S., Muley, S., LaConte, S. & Rottenberg, D. (2002), ‘The Quantitative Evaluation of

Functional Neuroimaging Experiments: The NPAIRS Data Analysis Framework', *NeuroImage* **15**(4), 747–771.
**URL:** *http://www.sciencedirect.com/science/article/pii/S1053811901910341*

Strother, S., La Conte, S., Hansen, L. K., Anderson, J., Zhang, J., Pulapura, S. & Rottenberg, D. (2004), 'Optimizing the fmri data-processing pipeline using prediction and reproducibility performance metrics: I. a preliminary group analysis', *Neuroimage* **23**, S196–S207.

Sutton, M., Thiébaut, R. & Liquet, B. (2018), 'Sparse partial least squares with group and subgroup structure', *Statistics in Medicine* **37**(23), 3338–3356.
**URL:** *https://onlinelibrary.wiley.com/doi/abs/10.1002/sim.7821*

Takane, Y. & Hwang, H. (2006), 'Regularized multiple correspondence analysis', *Multiple correspondence analysis and related methods* pp. 259–279.

Takane, Y. & Jung, S. (2009), 'Regularized nonsymmetric correspondence analysis', *Computational Statistics & Data Analysis* **53**(8), 3159–3170.
**URL:** *http://www.sciencedirect.com/science/article/pii/S0167947308004313*

Takane, Y., Yanai, H. & Mayekawa, S. (1991), 'Relationships among several methods of linearly constrained correspondence analysis', *Psychometrika* **56**(4), 667–684.

Tenenhaus, A., Philippe, C., Guillemot, V., Cao, K.-A. L., Grill, J. & Frouin, V. (2014), 'Variable selection for generalized canonical correlation analysis', *Biostatistics* p. kxu001.
**URL:** *http://biostatistics.oxfordjournals.org/content/early/2014/02/17/biostatistics.kxu001*

Tenenhaus, A. & Tenenhaus, M. (2011), 'Regularized Generalized Canonical Correlation Analysis', *Psychometrika* **76**(2), 257–284.
**URL:** *http://www.springerlink.com/content/8r17621w56k3025w/abstract/*

Tenenhaus, M. (1998), *La régression PLS: théorie et pratique*, Editions TECHNIP.

Tucker, L. R. (1958), 'An inter-battery method of factor analysis', *Psychometrika* **23**(2), 111–136.
**URL:** *http://www.springerlink.com/content/u74149nr2h0124l0/*

Wegelin, J. A. et al. (2000), 'A survey of partial least squares (pls) methods, with emphasis on the two-block case', *University of Washington, Tech. Rep* .

Winkler, A. M., Renaud, O., Smith, S. M. & Nichols, T. E. (2020), 'Permutation inference for canonical correlation analysis', *arXiv preprint arXiv:2002.10046* .

Wold, H. (1975), 'Soft modelling by latent variables: the non-linear iterative partial least squares (NIPALS) approach', *Journal of Applied Probability* **12**(S1), 117–142. Cambridge University Press.

Wold, S., Esbensen, K. & Geladi, P. (1987), 'Principal component analysis', *Chemometrics and Intelligent Laboratory Systems* **2**(1), 37–52.
**URL:** *http://www.sciencedirect.com/science/article/pii/0169743987800849*

Wold, S., Ruhe, A., Wold, H. & Dunn, III, W. (1984), 'The collinearity problem in linear regression. The partial least squares (PLS) approach to generalized inverses', *SIAM Journal on Scientific and Statistical Computing* **5**(3), 735–743.

Wold, S., Sjöström, M. & Eriksson, L. (2001), 'PLS-regression: a basic tool of chemometrics', *Chemometrics and Intelligent Laboratory Systems* **58**(2), 109–130.
**URL:** *http://www.sciencedirect.com/science/article/pii/S0169743901001551*
